# Reduced prediction error responses in high- as compared to low-uncertainty musical contexts

**DOI:** 10.1101/422949

**Authors:** D.R. Quiroga-Martinez, N.C. Hansen, A. Højlund, M. Pearce, E. Brattico, P. Vuust

## Abstract

Theories of predictive processing propose that prediction error responses are modulated by the certainty of the predictive model or *precision*. While there is some evidence for this phenomenon in the visual and, to a lesser extent, the auditory modality, little is known about whether it operates in the complex auditory contexts of daily life. Here, we examined how prediction error responses behave in a more complex and ecologically valid auditory context than those typically studied. We created musical tone sequences with different degrees of pitch uncertainty to manipulate the precision of participants’ auditory expectations. Magnetoencephalography was used to measure the magnetic counterpart of the mismatch negativity (MMNm) as a neural marker of prediction error in a multi-feature paradigm. Pitch, slide, intensity and timbre deviants were included. We compared high-entropy stimuli, consisting of a set of non-repetitive melodies, with low-entropy stimuli consisting of a simple, repetitive pitch pattern. Pitch entropy was quantitatively assessed with an information-theoretic model of auditory expectation. We found a reduction in pitch and slide MMNm amplitudes in the high-entropy as compared to the low-entropy context. No significant differences were found for intensity and timbre MMNm amplitudes. Furthermore, in a separate behavioral experiment investigating the detection of pitch deviants, similar decreases were found for accuracy measures in response to more fine-grained increases in pitch entropy. Our results are consistent with a precision modulation of auditory prediction error in a musical context, and suggest that this effect is specific to features that depend on the manipulated dimension—pitch information, in this case.

**Highlights:**

- The mismatch negativity (MMNm) is reduced in musical contexts with high pitch uncertainty
- The MMNm reduction is restricted to pitch-related features
- Accuracy during deviance detection is reduced in contexts with higher uncertainty
- The results suggest a feature-selective precision modulation of prediction error

Materials, data and scripts can be found in the Open Science Framework repository: http://bit.ly/music_entropy_MMN

DOI: 10.17605/OSF.IO/MY6TE

## 1. Introduction

Prediction is considered a core principle for brain function. Several theories propose that the brain anticipates and explains incoming information during perception based on its own predictive model of the world (Bar, 2009; Clark, 2016; Friston, 2005; Hohwy, 2013; Rao & Ballard, 1999). When unexpected information is encountered, it generates prediction error responses that are passed forward through brain hierarchies, so that they can drive perceptual inference, learning and action (den Ouden, Kok, & de Lange, 2012; Friston, 2010). Prediction error responses are hypothesised to depend, not only on the novelty of sensory signals, but also on the *precision* of the brain’s predictive model. Here we understand precision as the specificity and certainty of predictions, which can be driven both by the statistical properties of sensory signals and by internal factors such as attention. Precision is proposed to modulate the gain of prediction error responses so that they are stronger in perceptual contexts with low as compared to high uncertainty (Clark, 2013; Feldman & Friston, 2010; Hohwy, 2012). This mechanism, known as precision-weighting of prediction error, would ensure that primarily reliable perceptual contexts drive learning and behavior.

Research on predictive precision has mainly centered on the visual modality and selective attention as precision optimization (Feldman & Friston, 2010; Jiang, Summerfield, & Egner, 2013; Kok, Rahnev, Jehee, Lau, & de Lange, 2012). In the auditory domain, some studies have similarly focused on attention (Auksztulewicz & Friston, 2015; Chennu et al., 2013; Garrido, Rowe, Halász, & Mattingley, 2018; Schröger, Marzecová, & SanMiguel, 2015) and only a handful have shown how the statistical properties of the stimuli themselves can drive precision (Garrido, Sahani, & Dolan, 2013; Heilbron & Chait, 2018; Hsu, Bars, Hämäläinen, & Waszak, 2015; Sedley et al., 2016; Sohoglu & Chait, 2016; Southwell & Chait, 2018). These experiments, however, have employed very simple and artificial auditory stimuli, which limit the generality of the conclusions that can be drawn from them. As a result, it is not known how prediction error responses behave in more complex and realistic contexts. Consequently, the goal of the present study is to assess empirically whether prediction error responses are modulated by precision in a richer and more ecologically valid auditory context such as music.

Music perception provides a useful model of auditory prediction. Listeners are known to generate expectations in musical pieces, based on the statistical regularities of the context and long-term knowledge of a musical style (Huron, 2006; Pearce, 2018). The violation of these expectations generates neural prediction error responses (e.g. Carrus, Pearce, & Bhattacharya, 2013; Koelsch, Gunter, Friederici, & Schröger, 2000; Vuust et al., 2005). Interestingly, precision has been suggested to modulate musical prediction error and play an important role in the perceptual, aesthetic and emotional dimensions of musical experience (Hansen, Dietz, & Vuust, 2017; Ross & Hansen, 2016; Vuust, Witek, Dietz, & Kringelbach, 2018). Bringing empirical support to these claims, two behavioral studies have shown that listeners estimate the precision of musical expectations and that low-probability tones are judged as more unexpected in contexts with low as compared to high uncertainty (Hansen & Pearce, 2014; Hansen, Vuust, & Pearce, 2016). Nevertheless, how precision affects musical prediction error at the neural level remains unknown.

To address this question, we manipulated the precision of participants’ expectations by creating realistic melodic sequences with different degrees of pitch uncertainty. This was accomplished by manipulating two dimensions: the repetitiveness and the pitch alphabet—i.e. the collection of possible pitch categories—of the sequences. Research has shown that both factors modulate the neural signatures of predictive uncertainty (Auksztulewicz et al., 2017; Barascud, Pearce, Griffiths, Friston, & Chait, 2016). As a marker of prediction error, we recorded the mismatch negativity (MMN)(Näätänen, Gaillard, & Mäntysalo, 1978), which is a well-known neural response to regularity violations in a stimulus sequence (Näätänen, Paavilainen, Rinne, & Alho, 2007) with generators in the auditory and inferior frontal cortices (Deouell, 2007). The MMN is taken to reflect the violation and update of neural predictive models (Bendixen, SanMiguel, & Schröger, 2012; Garrido, Kilner, Stephan, & Friston, 2009; Lieder, Stephan, Daunizeau, Garrido, & Friston, 2013). Some studies already hint at a precision modulation of the MMN. In them, repetitive patterns are compared with random tone sequences that prevent the formation of regularities (Hsu et al., 2015; Jacobsen & Schröger, 2001; Maess, Jacobsen, Schröger, & Friederici, 2007), revealing no MMN for the latter. In other words, a very imprecise predictive model seems to lead to highly reduced prediction error responses. Perhaps for this reason, most MMN studies employ very simple and repetitive stimuli, which favor the strength of the recorded signal, but fail to provide a full picture of predictive processing in the rich and complex auditory environments of daily life. This is the case even for musical MMN paradigms that aim at making auditory stimuli more real-sounding (e.g., Tervaniemi, Huotilainen, & Brattico, 2014; Vuust et al., 2011). Therefore, employing stimuli that are more complex than in current paradigms, but at the same time less complex and more real-sounding than a random succession of tones, could reveal how predictive processing and precision operate in more ecologically valid settings.

The creation of more complex and realistic MMN paradigms comes with the challenge of establishing regularities in constantly changing acoustic streams. However, the auditory system is capable of extracting abstract features and complex relationships between sounds which give rise to an MMN when violated (Paavilainen, 2013). For example, in the no-standard multifeature paradigm (Pakarinen, Huotilainen, & Näätänen, 2010) deviants for different features immediately follow each other and regularities are created by keeping one feature constant for a period of time while tones change in other features. This suggests that features are processed independently from each other to a good extent, which is the property that we exploited here. In a novel MMN paradigm, we included intensity, timbre, pitch and slide deviants, and created regularities by keeping the same loudness, timbre, musical scale system and pitch steadiness—as opposed to pitch glide—in the sequence, even though standard tones constantly changed. Pitch deviants are of particular interest, since there was not a unique pitch height that served as standard. Rather, the regularity was created by the abstract properties of musical scales (Brattico, Tervaniemi, Näätänen, & Peretz, 2006), which constitute finite sets of possible pitch categories and distances between consecutive tones (i.e. intervals). Therefore, our pitch deviants consisted of out-of-tune sounds that fall outside the musical scale of reference (see section 2.1.2. for more details). Furthermore, the described feature independence is interesting because we manipulated precision only in the pitch dimension—by changing the number of possible pitch heights and intervals and their repetitiveness—while features such as intensity and timbre kept the same simple predictive model, with only one deviant and one standard value across conditions. In consequence, our multifeature paradigm provided an opportunity to explore the extent to which precision in one auditory dimension affects predictive processing in other auditory dimensions.

In the present study, we conducted separate neurophysiological and behavioral experiments to determine whether auditory prediction error is modulated by precision during listening to musical sequences. In the neurophysiological experiment, we used magnetoencephalography (MEG) to record magnetic mismatch responses (MMNm) to pitch, intensity, timbre and slide deviants in high-entropy (HE) and low-entropy (LE) contexts. The LE context consisted of an adapted version of the simple and repetitive musical multi-feature paradigm (Vuust et al., 2011) (see section 2.1.2. for more details), whereas the HE context consisted of a set of novel non-repetitive melodies. Entropy was quantitatively characterized with a computational model of auditory expectation (see section 2.1.2.1). In line with a precision modulation of prediction error, we expected reduced—but still present—MMNm amplitudes for the HE as compared to the LE context. Moreover, in exploratory analyses we addressed the possibility that different features were affected by pitch entropy in different ways, thereby providing a first investigation of the feature-selectivity of the effect. In the behavioral experiment, we asked participants to detect pitch deviants introduced in several tone sequences, and to report the confidence of their responses. This experiment had two aims. The first was to assess behaviorally the putative precision modulation of the MMNm. The second was to determine whether listeners are sensitive to fine-grained manipulations of precision. For this reason, we employed stimuli with five degrees of entropy, which included a subset of the HE/LE stimuli used in the MEG experiment and three additional conditions with intermediate entropy levels. We expected lower accuracy and confidence ratings as the entropy of the melodic sequences increased.

## 2. Materials and Methods

The data, code and materials necessary to reproduce the reported experiments and results are available at http://bit.ly/music_entropy_MMN; DOI 10.17605/OSF.IO/MY6TE.

### 2.1. MEG experiment

#### 2.1.1. Participants

Twenty-four right-handed and neurologically healthy non-musicians (13 women, mean age 27, range 19-34) took part in the experiment. The sample size was chosen to be similar to a previous study using the same MEG scanner in which relatively small within-subjects differences were identified (Hansen, Højlund, Møller, Pearce, & Vuust, 2019). Musical expertise was measured with the musical training subscale of the Goldsmiths Musical Sophistication Index (Gold-MSI) questionnaire, which has been validated in a very large sample (*n*⎕=⎕147,636) (Müllensiefen, Gingras, Musil, & Stewart, 2014). The mean score was 10.3 (SD = ± 3.5) and all scores lay in the 26^th^ percentile of the norm for the subscale. Moreover, participants’ musical competence was assessed with the Musical Ear Test (MET), which has been shown to accurately discriminate between musicians and nonmusicians (Wallentin, Nielsen, Friis-Olivarius, Vuust, & Vuust, 2010). The test yielded a total score of 69.12 (SD = ± 9.58), which falls within normal values for this population (Wallentin et al., 2010). Participants were recruited through an online database for experiment participation, agreed to take part voluntarily, gave their informed consent and received 300 Danish kroner (approximately 40 euro) as compensation. The data from all participants were included in the analyses, since reliable auditory responses were identified in all cases, which was our predefined inclusion criterion. The study was conducted in accordance with the Helsinki declaration and was approved by the regional ethics committee (De Videnskabsetiske Komitéer for Region Midtjylland in Denmark).

#### 2.1.2. Stimuli

High-entropy and low-entropy stimuli were included (Figure 1a). For the LE condition, we adapted the original musical multi-feature paradigm (Vuust et al., 2011; Vuust, Brattico, Seppänen, Näätänen, & Tervaniemi, 2012; Vuust, Liikala, Näätänen, Brattico, & Brattico, 2016), which consists of a four-note repeating pattern (low-high-medium-high pitch) employing the notes from a major or minor chord. This pattern, known as the Alberti bass, is used across musical styles (Fuller, 2001). The HE condition consisted of a novel multi-feature paradigm including a set of six novel melodies which contained almost no exact repetitions of pitch patterns and a larger pitch alphabet than LE. All the tones in the melodies were isochronous to make both conditions directly comparable with each other and other MMN paradigms (the full set of stimuli is shown in supplementary file 1). Individual tone sequences—i.e., single melodies or Alberti bass sequences—in both conditions were 32-notes long, lasted eight seconds and were pseudo-randomly transposed from 0 to 5 semitones upwards to the keys comprising the major and minor modes of C, C#, D, D#, E, and F. After transposition, the pitch alphabet in the HE condition spanned up to thirty-one semitones from B3 (F_0_ ≈ 247 *Hz*) to F6 (F_0_ ≈ 1397 *Hz*). To minimize acoustic confounds, we made sure that LE sequences spanned approximately the same global pitch alphabet as HE sequences by transposing half of them to the octave from C4 (F_0_ ≈ 262 *Hz*) to C5, and the other half to the octave from C5 to C6 (supplementary figure 1).

**Figure 1.**
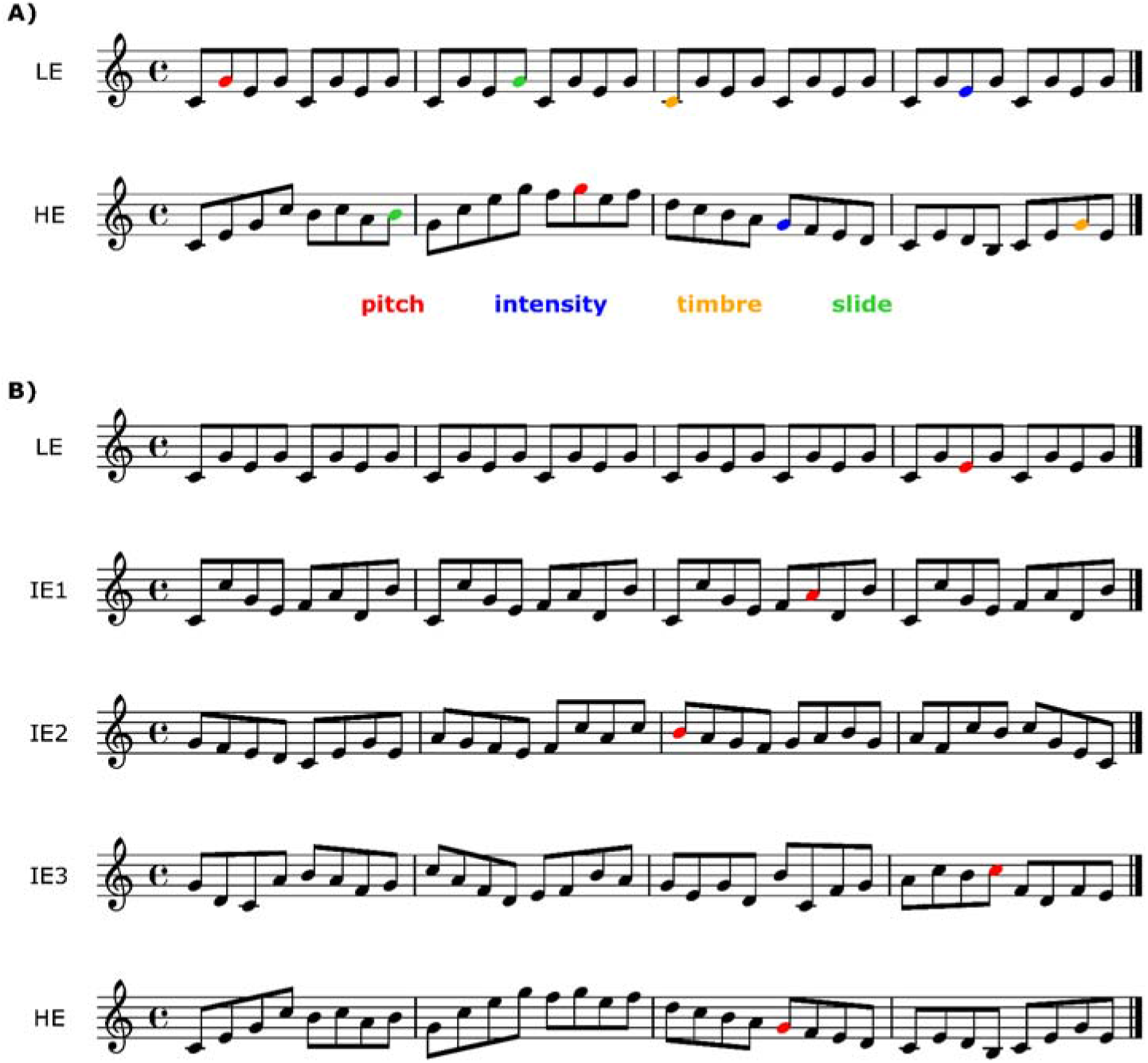
Examples of the sequences used in A) the MEG experiment and B) the behavioral experiment (complete stimulus set available in the supplementary file 1 and sound examples available in the online repository). LE = low entropy, IE = intermediate entropy, HE = high entropy. Two of the conditions in the behavioral experiment (LE, HE) correspond to the conditions in the MEG experiment.

Stimuli were presented using a pool of piano tones made with the “Warm Grand” sample from the Halion sampler in Cubase (Steinberg Media Technology, version 8). Each tone had a duration of 250 ms, was peak-amplitude normalized and had 3-ms-long fade-in and fade-out to prevent clicking. This tone duration shortened stimulation time while preventing the MMN from overlapping with the onset of the following tone. No gaps were introduced between consecutive sounds to create the perception of continuous melodic phrases with legato articulation. We regard this as representative of how melodies are played in real music. Pitch deviants consisted of out-of-tune tones falling outside the musical scale of reference and were created by raising the pitch of the original tones by 50 cents. The slide deviant was a continuous pitch glide which spanned the whole duration of the tone, going from two semitones below towards the pitch of the corresponding standard tone. For the intensity deviant sound level was decreased by 20 dB. The timbre deviant consisted of a telephone receiver effect (bandpass-filtered between 1 and 4 kHz). All deviants were created with Audition (Adobe Systems Incorporated, version 8).

Each condition was presented in a separate group of three consecutive blocks. Within each block, tone sequences were played one after the other without pauses. The order of HE sequences was pseudorandom so that any sequence of twelve consecutive melodies contained no more than one major and minor version of each. No melody was played twice in a row. Transpositions in both conditions were pseudorandomized in the same way. At the beginning of each block, a sequence with no deviants was added to ensure a certain level of auditory regularity at the outset. The duration of the pause between blocks was not fixed but usually took around one minute.

Deviants were introduced as follows. Each 32-note sequence was divided into eight groups of four notes (Figure 1a). In half of the sequences, deviants occurred in groups 1, 3, 5 and 7. In the other half, they occurred in groups 2, 4, 6 and 8. This was done because we also included a combined condition where HE and LE sequences were played simultaneously, thereby creating two-part musical excerpts. Thus, the position of the deviants was distributed across streams to counterbalance the effects of key-changes between parts. The purpose of this condition was to assess the predictive processing of simultaneous musical streams, which is beyond the scope of this article. The corresponding results will be reported elsewhere. Within each four-note group, only one deviant could occur randomly in any of the four positions with equal probability. There was one deviant per feature in each sequence and their order of appearance was pseudorandom. There were 144 sequences in each condition and the same number of deviants per feature. This number was close to the minimum of 150 suggested by Duncan et al. (2009) for EEG research in clinical populations. Since each deviant type occurred once per thirty-two notes, its overall zeroth-order probability was 1/32 ≈ 0.031. In the session, we also included another group of three consecutive blocks in which Alberti bass sequences were played in a low pitch range. This condition served as a control for the combined condition and therefore is not the focus of this article either. The order of this and the HE and LE conditions was counterbalanced across participants. These conditions always came after the three blocks of the combined condition.

##### 2.1.2.1. Quantitative estimates with IDyOM

To characterize quantitatively the stimuli, we used Information Dynamics of Music (IDyOM), a variable-order Markov model of expectation (Pearce, 2005, 2018). IDyOM generates expectations at each point of an event sequence in the form of a probability distribution (*P*) over the set of possible continuations at that particular moment. These probabilities are conditional on the preceding context and the previous long-term exposure of the model. The uncertainty of expectations is quantified in terms of Shannon entropy:

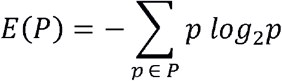

Since the probabilities of the possible continuations (*p*) sum to one, entropy is minimal when only one event has a very high probability, and is maximal when all possible events are equally likely. IDyOM’s entropy estimates correlate with behavioral measures of uncertainty (Hansen & Pearce, 2014; Hansen et al., 2016). Once the next event in the sequence appears, IDyOM estimates its unexpectedness as information content (IC):

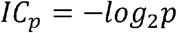

Thus, unexpected or surprising events have high IC. Here, mean entropy and mean IC values are used to estimate how the statistical properties of the sequences drive listeners’ predictive precision (see section 2.1.2.1. for further details). Our primary measure in this context is entropy, as it directly estimates precision, given that precision is the inverse of uncertainty. Thus, contexts with low entropy would generate more precise expectations than contexts with high entropy. Our secondary measure is mean IC, since stimuli with decreasing levels of precision—and thus increasing levels of entropy— would tend to yield higher levels of unexpectedness and IC in the long run. Here we consider both measures to obtain a more complete picture of the listener’s predictive model in relation to the stimuli. Note, however, that our manipulations are qualitative, and that we use IDyOM merely to characterize the uncertainty of the previously generated stimuli rather than to directly predict neural activity.

Mean entropy and IC values were quantified using a model that combined short-term probabilities inferred from the sequences themselves with long-term probabilities learned from a corpus of Western tonal hymns and folksongs (datasets 1, 2, and 9 from Table 4.1 in Pearce 2005, comprising 50,867 notes). This corpus has been extensively used in prior research. The model simulates a listener that generates predictions based on life-long knowledge of Western tonal music, but who is also capable of learning and incorporating the structure of the current stimuli into its long-term expectations. This configuration is known as *Both+* model. IDyOM can use different parameters of the musical surface—known as viewpoints—to derive its probabilistic predictions (Conklin & Witten, 1995; Pearce, 2005). Research has often used a viewpoint combining tonal scale degree (i.e. the perceived stability of a given pitch with respect to its tonal context) and pitch-interval (e.g., Carrus et al., 2013; Hansen & Pearce, 2014; Omigie, Pearce, Williamson, & Stewart, 2013; Pearce, Ruiz, Kapasi, Wiggins, & Bhattacharya, 2010) in order to capture both melodic and tonal structure (see Pearce, 2005, for more details). This viewpoint was used here to obtain note-by-note IC and entropy values, which were then averaged for each sequence and condition. Crucially, IDyOM uses the pitch alphabet of the training corpus for its predictions, which in this case was larger than the alphabets of our stimuli. This produces misleading estimates and makes the model insensitive to differences in pitch alphabet between conditions, which have been identified as an important source of uncertainty (Auksztulewicz et al., 2017; Barascud et al., 2016). For this reason, we adjusted the distributions to include only the probabilities of the tones present in each condition, which were then renormalized to sum to one (for non-adjusted values see supplementary figure 3). The analysis revealed higher mean entropy and IC for HE than LE stimuli (Figures 2a and 2c). IC and entropy profiles of all the sequences are shown in supplementary file 1.

**Figure 2.**
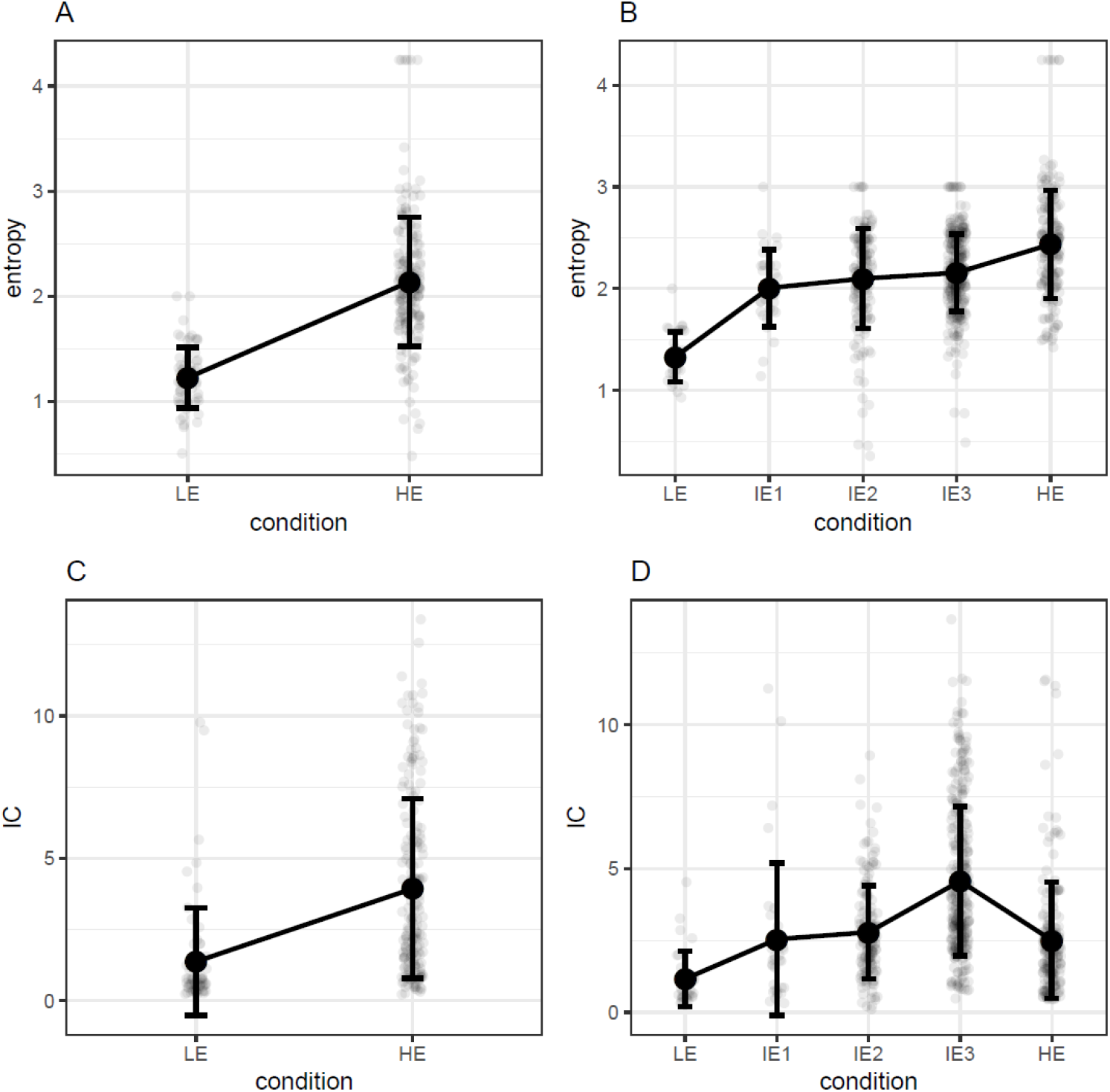
Entropy and information content (IC) values, in bits, measured with IDyOM for the stimuli included in the MEG (A, C) and behavioral (B, D) experiments. Error bars represent standard deviation. LE= Low entropy, IE = Intermediate entropy, HE = High entropy.

#### 2.1.3. Procedures

Participants received oral and written information and gave their consent. Then they filled out the Gold-MSI questionnaire and completed the MET. Once participants had put on MEG-compatible clothing, electrodes and coils were attached to their skin and their heads were digitized. During the recording, they were sitting upright in the MEG device looking at a screen. Before presenting the musical stimuli, their auditory threshold was measured through a staircase procedure and the sound level was set at 60dB above threshold. Participants were instructed to watch a silent movie of their choice, ignore the sounds and move as little as possible. This task minimizes the overlap of the MMN with attention-related components such as the P300 (Duncan et al., 2009). Participants were told there would be musical sequences playing in the background interrupted by short pauses so that they could take a break and readjust their posture. Sounds were presented through isolated MEG-compatible ear tubes (Etymotic ER•30). The recording lasted approximately 90 minutes and the whole experimental session took between 2.5 and 3 hours including consent, musical expertise tests, preparation, instructions, breaks, and debriefing.

#### 2.1.4. MEG recording and analyses

Magnetic correlates of brain activity were recorded using an Elekta Neuromag MEG TRIUX system with 306 channels (204 planar gradiometers and 102 magnetometers) and a sampling rate of 1000 Hz. Participants’ head position was monitored with four coils (cHPI) attached to the forehead and the mastoids. Offline, signals coming from inside the skull were isolated with the temporal extension of the signal source separation (tSSS) technique (Taulu, Kajola, & Simola, 2003) using Elekta’s MaxFilter software (Version 2.2.15). This procedure included movement compensation for all but two participants, for whom continuous head position information was not reliable due to suboptimal placement of the coils. In these cases, the presence of reliable auditory event-related fields (ERFs) was successfully verified by visually inspecting the amplitude and polarity of the P50(m) component. Eye-blink and heartbeat artifacts were corrected with the aid of electrocardiography, electrooculography and independent component analysis, as implemented by a semi-automatic routine (FastICA algorithm and functions find_bads_eog and find_bads_ecg in the software MNE-Python) (Gramfort, 2013). Visual inspection served as quality check.

The ensuing analysis steps were conducted with the Fieldtrip toolbox (version r9093) (Oostenveld, Fries, Maris, & Schoffelen, 2011) in MATLAB (R2016a, The MathWorks Inc., Natick, MA). Epochs comprising a time window of 400 ms after sound onset were extracted and baseline-corrected, with a pre-stimulus baseline of 100 ms. Epochs were then low-pass filtered with a cut-off frequency of 35 Hz and down-sampled to a resolution of 256 Hz. For each participant, ERFs were computed by averaging the responses for all deviants for each feature and averaging a selection of an equal number of standards. These were selected by finding, for each single deviant, a standard tone that was not preceded by a deviant and was in the same position of the same HE or LE sequence—although not necessarily the same transposition—in a different trial. This ruled out artefacts related to the difference in noise between conditions—since there are many more standards than deviants—and the position of the deviant within the sequence. After averaging, planar gradiometers were combined by computing root mean square values. Finally, a new baseline correction was applied and MMNm difference waves were computed by subtracting the ERFs of standards from the ERFs of deviants.

The statistical analyses were performed on combined gradiometer data, as these sensors measure activity directly above the neural sources and have a better signal-to-noise ratio (Haumann, Parkkonen, Kliuchko, Vuust, & Brattico, 2016). Magnetometers were used to inspect the polarity of components. For the primary analyses, we used two-sided paired-samples *t*-tests in a univariate-mass approach with cluster-based permutations as multiple comparisons correction (Maris & Oostenveld, 2007). The sample-level significance threshold was .05, the chosen statistic was the maximal sum of clustered *T*-values (*maxsum*) and the number of iterations was 10,000. The tests were restricted to a time window between 100 and 250 ms after sound onset as this covers the typical latency of the MMNm (Näätänen et al., 2007). To assess the presence of MMNm responses, the ERFs of deviants and standards were contrasted for each feature and condition. To evaluate the effect of stimulus entropy, the MMNm difference waves in the HE and LE conditions were contrasted for each feature. Since separate tests were performed for each feature, a Bonferroni correction for multiple comparisons was applied by multiplying *p*-values by the number of features, namely four.

Further exploratory analyses on mean gradient amplitudes (MGA) were performed to estimate whether MMNm responses for different features were affected differently by stimulus entropy. These analyses were not conducted with a univariate-mass approach since peak latencies were clearly different between features, which does not allow a direct comparison of amplitude. Instead, MGAs were computed for each participant, feature and condition, by averaging ±25 ms around the peak, defined as the highest local maxima of the MMNm difference wave between 100 and 250 ms. This procedure was restricted to the average of the four combined gradiometers in each hemisphere with the largest P50(m) responses in the grand average, which we regard as an indicator of reliable auditory signals (channels in red in the top-right head-plot of Figure 3). These channels also exhibit the largest MMNm amplitudes (Figure 4). Differences in MGA between conditions were computed and used as the dependent variable. A linear mixed model including feature, hemisphere and their interaction as predictors was fitted. Random intercepts were also included. Random slopes were excluded as the amount of data points was not sufficient for their estimation. Since this analysis was exploratory, we report parameter estimates and confidence intervals, but not *p*-values. Standardized effect sizes (Cohen’s *d*) of pairwise comparisons were also computed as the difference between means divided by the residual standard deviation.

**Figure 3.**
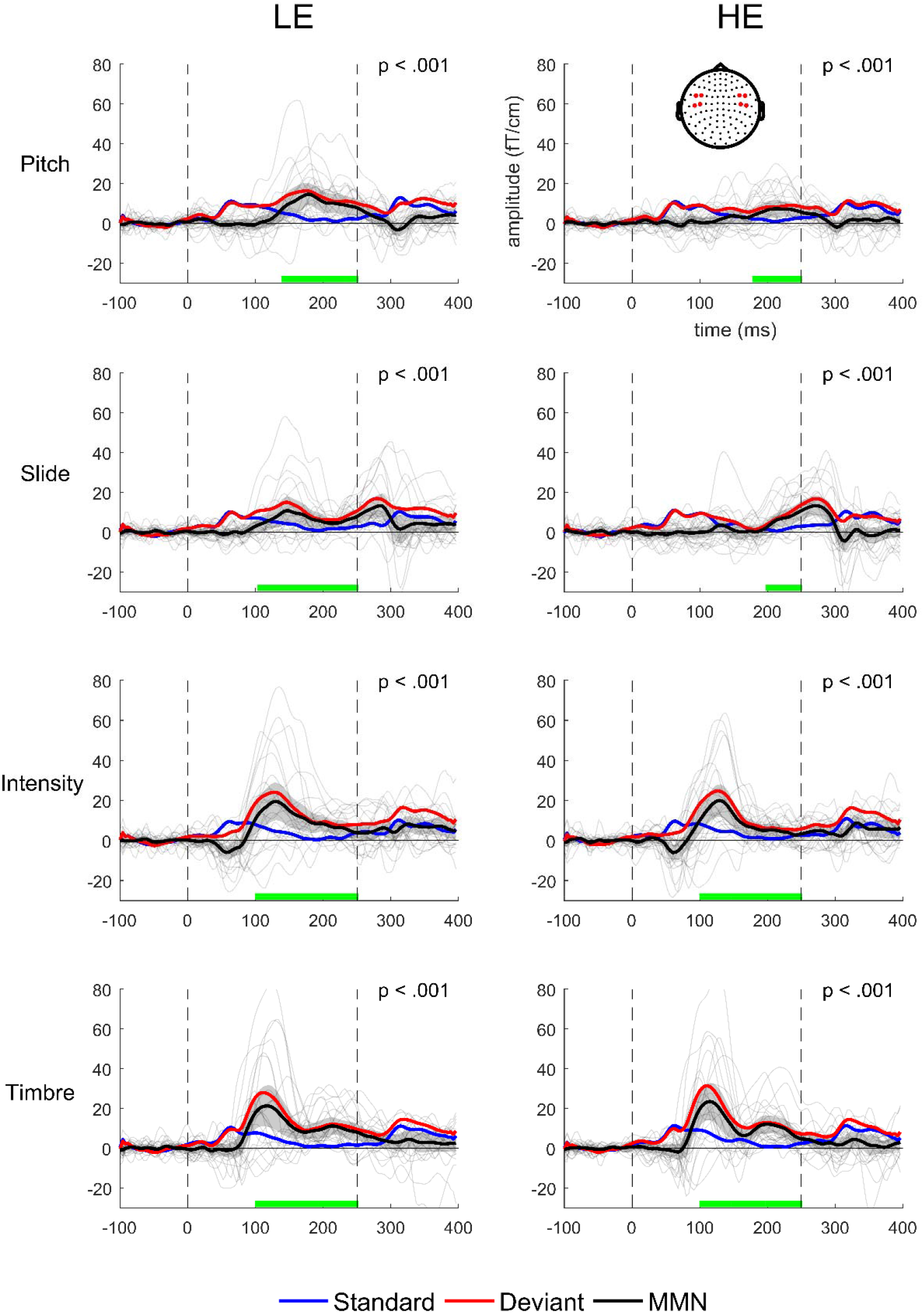
Standard, deviant and difference (MMNm) amplitudes in high- (HE) and low-entropy (LE) conditions. Traces correspond to the average of the four right temporal combined gradiometers shown in red in the top-right head plot. Gray lines show individual MMNm traces. Dashed vertical lines indicate sound onsets. Shaded areas indicate 95% confidence intervals. For descriptive purposes, green lines indicate the time where the difference between standards and deviants was significant. Note, however, that this interval is not a reliable estimate of the true extent of the effect (Sassenhagen & Draschkow, 2019).

### 2.2. Behavioral experiment

In this experiment, a deviance detection task was used to confirm behaviorally the hypothesized neurophysiological effects. In addition, since the MEG experiment included only two highly contrasting conditions to observe clear differences in the neural signal, in the behavioral experiment we aimed to assess a more fine-grained precision modulation of prediction error, and test whether repetitiveness or pitch alphabet alone are sufficient to elicit the effect.

#### 2.2.1. Participants

Twenty-one non-musicians (16 women, mean age 21.9, range 18-36) participated in the experiment. Musical expertise was measured with the Gold-MSI musical training subscale which yielded a score of 12.9 (SD = ±5.77). All values lay in the 42^nd^ percentile of the norm for this subscale. Participants were recruited through an online database for experiment participation, agreed to take part voluntarily, gave their informed consent and received 100 Danish kroner (approximately 13.5 euro) as compensation. Two subjects had previously participated in the MEG experiment. The data from all participants were analyzed, since above-chance deviance detection was verified in all cases. This was our predefined inclusion criterion. The sample size was chosen to be comparable to that of the MEG experiment. In this regard, Bishop and Hardiman (2010) demonstrated that behavioral deviance detection is present even for subjects who do not show reliable individual MMN responses, thus suggesting a higher sensitivity of behavioral measures.

#### 2.2.2. Experimental design

Participants were presented with 32-note sequences with different levels of entropy and asked to decide after each one if a note with a wrong pitch was present in the sequence or not, and how certain they were about their response on a scale from 1 (not certain at all) to 7 (completely certain). Five conditions with different degrees of entropy were included (Figure 1b; the full stimulus set is shown in supplementary file 1). As in the MEG experiment, there was an LE condition consisting of an Alberti bass sequence, and an HE condition corresponding to a subset of five of the six melodies used in the MEG session. Three intermediate conditions (IE) were added to test for more fine-grained effects of entropy. The alphabet of these conditions was restricted to a C-major scale spanning eight tones from C4 to C5. Based on previous research showing an effect of pitch alphabet on the uncertainty of auditory stimuli (Auksztulewicz et al., 2017; Barascud et al., 2016), we conjectured that these sequences would have higher mean entropy than LE sequences, which spanned only three pitch categories, and lower mean entropy than HE sequences, which spanned up to fifteen pitch categories. For the three intermediate conditions, entropy was manipulated by changing their repetitiveness. Thus, the least uncertain of the three (IE1) was a repeated eight-note pattern. The middle condition (IE2), which consisted of five melodies, relaxed the constraint for exact repetition leading to reduced precision over the IE1 condition. Finally, the most uncertain of the three conditions (IE3) consisted of random orderings of the eight tones, with equal probability and without playing any of them twice in a row. These sequences were generated individually for each participant. Since sequential constraints are minimal in this condition, it was expected to have higher entropy than IE1 and IE2, but lower entropy than HE, given its smaller pitch alphabet. Note that the contrast LE-IE1 would reveal whether pitch alphabet is sufficient to elicit an uncertainty effect whereas the contrasts IE1-IE2, IE1-IE3 and IE2-IE3 would show the same with regard to repetitiveness.

The conjectured pattern (LE < IE1 < IE2 < IE3 < HE) was confirmed by IDyOM’s mean entropy values (Figure 2b), estimated as described in section 2.1.2.1. Mean IC values followed a similar pattern (LE < HE < IE1 < IE2 < IE3), with the exception that HE had lower IC than all the intermediate conditions (Figure 2d). This might reflect the fact that HE melodies tended to have smaller pitch intervals (mostly 1- or 2-semitone steps) which are more common than larger intervals in Western tonal music (Huron, 2006) (supplementary figure 2). These estimates were not meant to be used as predictors in the analyses. Rather, they were used as approximate values to confirm the putative ordering of the conditions and help the interpretation of the results. Note that, since IE3 (random) sequences were unique for each subject, in this case entropy and IC values were estimated only for one participant, as a representative sample. These are the ten sequences shown in supplementary file 1. Finally, even though the variance of single-tone estimates was high, mean values for individual melodies showed little overlap between conditions (supplementary figure 4), which indicates that entropy and IC systematicaly changed for each sequence according to our manipulations. This means that any effects observed in this study would be most likely driven by context entropy because, if uncertainty were mainly driven by single-note entropy, the high variance would add a lot of noise to behavioral measures and potentially make the effects undetectable.

To simplify the stimuli and make them comparable between conditions, only sequences in the C-major key were included. Target sequences were created by randomly choosing a tone from the second half of each sequence and raising its pitch 25 cents up. Deviations were smaller than in the MEG experiment to avoid ceiling effects observed during piloting. Only pitch deviants were included, since this feature showed the strongest reduction in amplitude between conditions in the neurophysiological data (see section 3.1.2). The creation of the pool of standard and deviant tones followed the same procedure as in the MEG experiment. Ten targets and ten foil sequences were presented for each condition in a random order. We chose the number of trials as a compromise between the length of the task, the number of conditions and the amount of data required to produce reliable estimates, based on pilot tests. Note that the number of possible sequences differed between conditions which meant that they were repeated a different number of times. Thus, LE and IE1 consisted of only one sequence, and for this reason they were repeated ten times as targets and ten times as foils. In contrast, IE2 and HE consisted of five different sequences, which entailed that they were repeated twice as targets and twice as foils. Finally, since there were ten unique IE3 sequences for each participant, they were played only once as foils and once as targets. There were four practice trials at the beginning of the session. The complete procedure lasted approximately 25 minutes.

#### 2.2.3. Statistical analyses

We used signal detection theory to analyze deviance detection performance (Stanislaw & Todorov, 1999). Both *d’*- and criterion (*c*-) scores were computed for each participant and condition. In the few cases where a participant scored 100% or 0% of hits or false alarms, values were adjusted to 95% or 5%, respectively. This prevented the *z* values in the computations from reaching infinity. *d’*-scores quantify the difference between the proportions of hits and false alarms. Therefore, they provide a more accurate measure than hit rates, since they take into account the bias in the response—e.g. answering always yes. This bias can be directly quantified by *c*-scores, measured as the negative average of the proportion of hits and false alarms.

Statistical analyses were performed using the software *R* (R Core Team, 2019). To assess the effects of stimulus entropy, mixed models were fitted using the *lmer* function from the *lme4* package (Bates, Mächler, Bolker, & Walker, 2015). The models allowed a random intercept for each participant to account for individual differences. No random slopes were included, since there were not enough data points to avoid overfitting and reach convergence. Three different models were compared: one with an intercept only (*d0*), another with an additional term for entropy as a categorical factor (*d1*), and a final model (*d2*) with the five conditions treated as an ordered linear predictor, asigning values from 1 to 5 according to our entropy manipulations. The comparison between *d1* and *d0* assessed the overall effect of entropy, whereas comparing *d2* and *d1* revealed to what extent a linear trend could explain the data. Regarding *c*-scores, we compared an intercept-only model (*cr0*) with a model including entropy as a categorical factor (*cr1*), to assess the extent to which bias changed between conditions. Akaike Information Criteria (AIC) and likelihood ratio tests were used for all comparisons.

Regarding confidence scores, ordinal logistic regression was employed in the form of a cumulative link mixed model (Christensen, 2015), as implemented by the function clmm from the *ordinal* package (Christensen, 2018). Log-odds (“logit”) was the link function. This method allowed the quantification of the change in the proportion of responses in each confidence category, relative to the entropy conditions. We fitted three initial models (*c0, c1, c2*) in which the estimated parameters, random effects and model comparisons were the same as in the analysis of d’-scores, with the only difference that now there was an intercept for each of the six cut-points between response categories (see suplementary file 2). Moreover, we fitted two additional models (*c1s, c2s*) including random slopes for the effect of entropy (categorical and ordered, respectively), since the amount of data made it possible in this case. Finally, post-hoc, Bonferroni-corrected pairwise comparisons between conditions were conducted with the function *glht* (from the *multcomp* package, Hothorn, Bretz, & Westfall, 2008) for *d’*- and *c*-scores, and *lsmeans* (from the *lsmeans* package, Lenth, 2016) for confidence ratings.

## 3. Results

### 3.1. MEG experiment

#### 3.1.1. Presence of the MMNm

Significant differences were found between standard and deviant ERFs in the 100-250 ms post-stimulus time window for all features in both conditions (Figure 3). The differences were present bilaterally, were largest over right temporal gradiometers, and showed a polarity—as observed in the magnetometers—consistent with previous reports of the MMNm (e.g., Bonetti, Haumann, Vuust, Kliuchko, & Brattico, 2017) (Figure 4).

**Figure 4.**
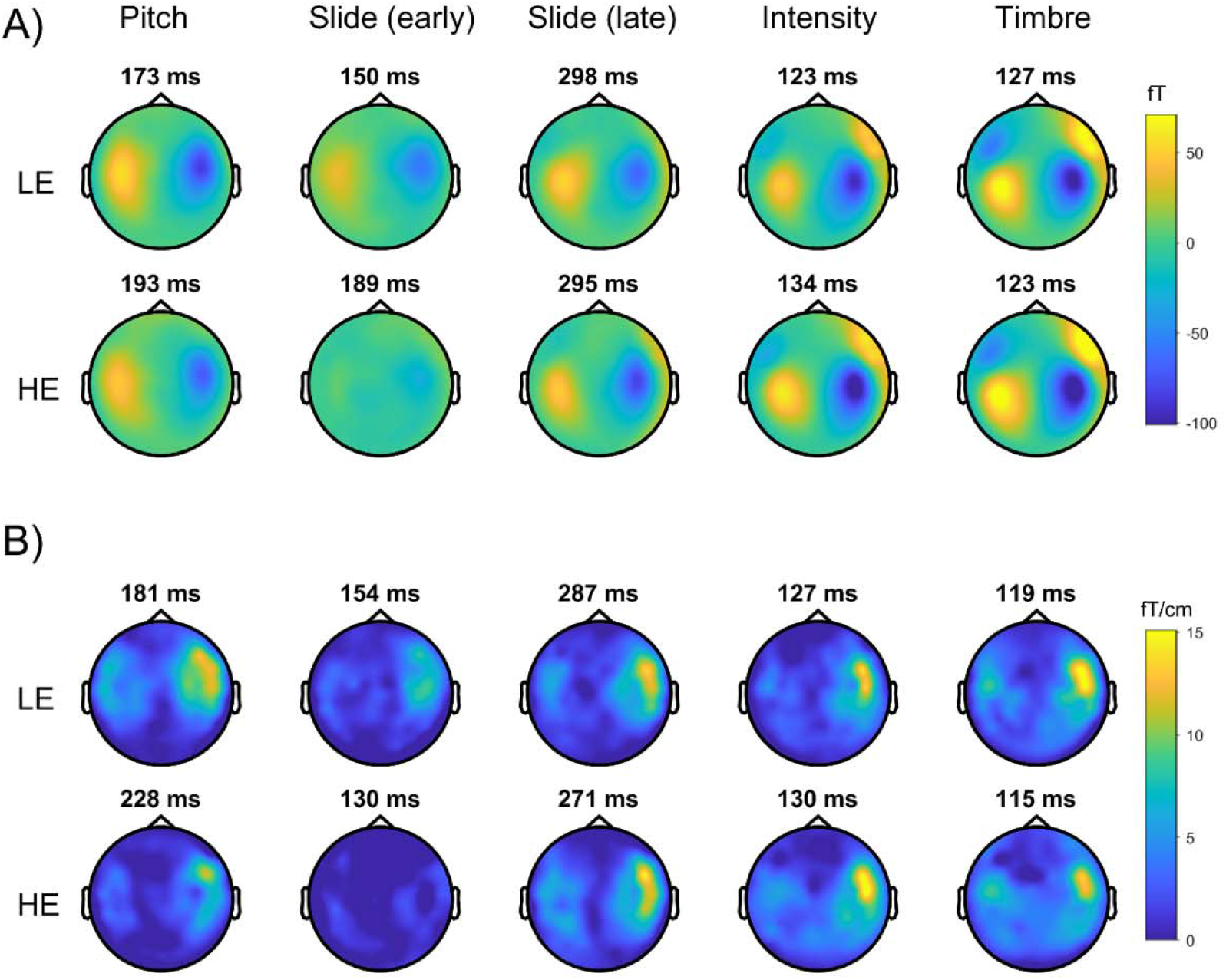
Topography of the MMNm (difference between standards and deviants) for (A) magnetometers and (B) gradiometers. Peak latencies are shown above each plot. The displayed activity corresponds to an average of *±*25 ms around the peak. Slide MMNm topographies are shown for activity around the peak in both early (100-250 ms) and late (250-350 ms) time windows. LE = low entropy, HE = high entropy

#### 3.1.2. Low-entropy vs. high-entropy stimuli

An amplitude reduction in the MMNm difference waves was found for HE as compared to LE stimuli, for pitch and slide deviants bilaterally (Figure 5). This reduction was maximal at temporal gradiometers. No significant differences were found for intensity or timbre. The exploratory MGA analyses suggested that differences between HE and LE contexts were larger for pitch and slide as compared to timbre and intensity, in the right hemisphere (*d > .6*) (Figure 6a, Table 1). From the pairwise comparisons, only the ones between pitch and timbre and slide and timbre yielded a 95% confidence interval excluding zero. Differences in the left hemisphere, as well as between pitch and slide or intensity and timbre, were small (*d* ≤ .4).

**Figure 5.**
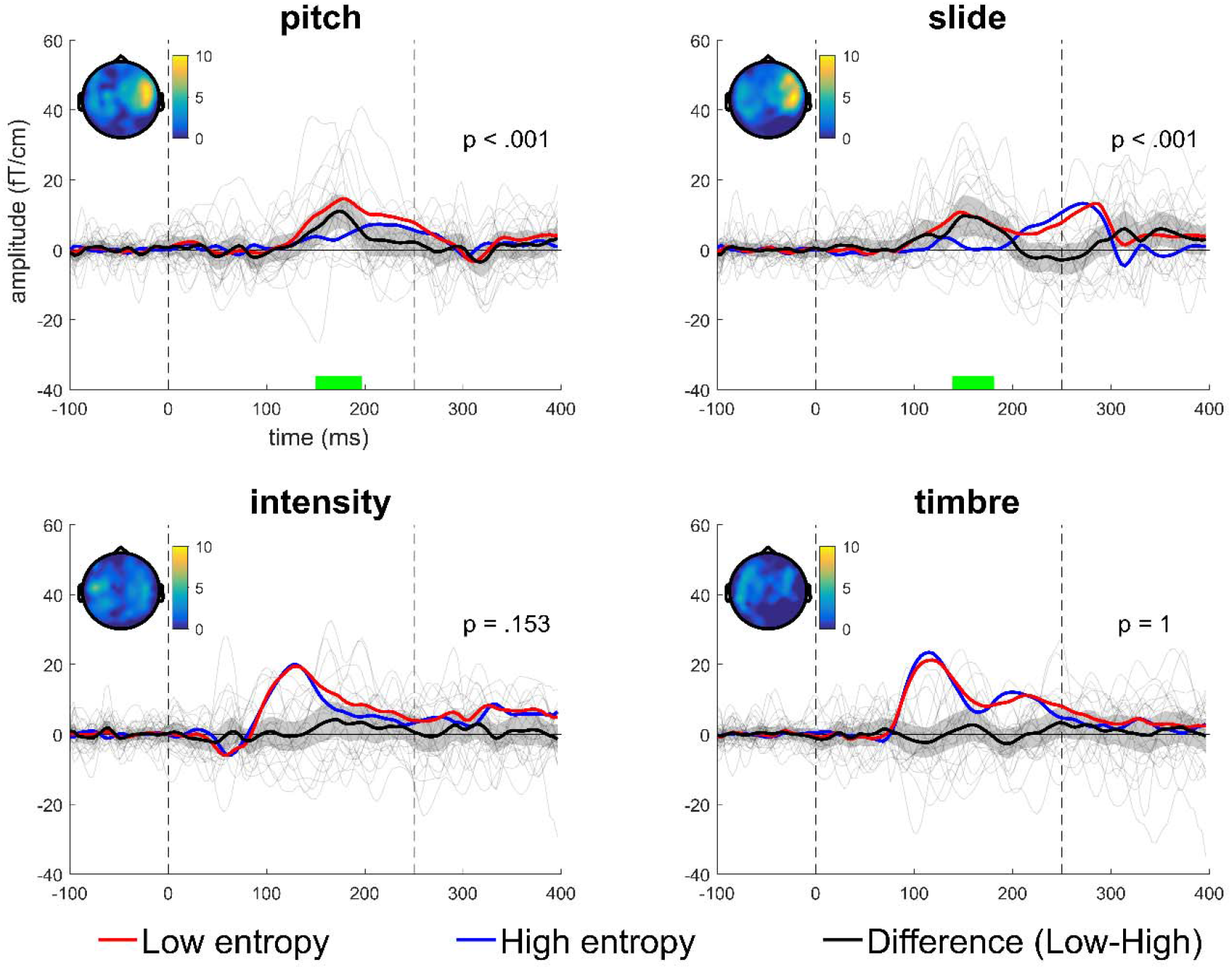
MMNm difference waves according to the two conditions. Traces correspond to the average of the four right temporal combined gradiometers shown in red in the top-right head plot on Figure 3. Gray lines show individual differences. Dashed vertical lines indicate sound onsets. Shaded areas indicate 95% confidence intervals. Topomaps show activity *±*25 ms around the peak difference. For descriptive purposes, green lines indicate the time where the difference between conditions was significant. Note, however, that this interval is not a reliable estimate of the true extent of the effect (Sassenhagen & Draschkow, 2019).

**Table 1.** Pairwise contrasts between features for the MMNm amplitude differences between low-entropy and high-entropy conditions. p = pitch, s = slide, i = intensity, t = timbre.

| contrast | hemisphere | estimate | CI 2.5% | CI 97.5% | t | Cohen's d |
| --- | --- | --- | --- | --- | --- | --- |
| p - s | right | 0.4 | -4.4 | 5.19 | 0.21 | 0.06 |
| p - i | right | 4.68 | -0.11 | 9.47 | 2.53 | 0.75 |
| p - t | right | 7.18 | 2.39 | 11.98 | 3.89 | 1.15 |
| s - i | right | 4.28 | -0.51 | 9.08 | 2.32 | 0.68 |
| s - t | right | 6.79 | 2 | 11.58 | 3.67 | 1.08 |
| i - t | right | 2.51 | -2.29 | 7.3 | 1.36 | 0.4 |
| p - s | left | -2.31 | -7.1 | 2.48 | -1.25 | -0.37 |
| p - i | left | -1.58 | -6.38 | 3.21 | -0.86 | -0.25 |
| p - t | left | 0.06 | -4.74 | 4.85 | 0.03 | 0.01 |
| s - i | left | 0.73 | -4.07 | 5.52 | 0.39 | 0.12 |
| s - t | left | 2.37 | -2.43 | 7.16 | 1.28 | 0.38 |
| i - t | left | 1.64 | -3.15 | 6.43 | 0.89 | 0.26 |

The slide MMNm displayed an unusual shape (Figures 3 and 5). Specifically, it was sustained beyond 250 ms and peaked around 280 ms. The magnetometer polarity of the late response was the same as that of the MMNm for the other features (Figure 4a). Furthermore, stimulus entropy seemed to affect the earlier portion of the ERF more than the later part. In an exploratory analysis, a mixed effects model with random intercepts, and latency and hemisphere as predictors, suggested larger differences in the earlier than the later time window and no substantial differences as a function of hemisphere or the interaction between hemisphere and latency (Figure 6b, parameters reported in the supplementary file 2).

### 3.2. Behavioral experiment

The analyses of *d*’-scores showed that the *d1* model (with a term for entropy) explained the data better than the *d0* (intercept-only) model (*χ^2^* = 39.31, *p* < .001). The AIC value was 269.43 for *d0* and 238.12 for *d1*, which agrees with the likelihood ratio test. Moreover, the comparison between *d1* and *d2* (a model with entropy as an ordered linear variable) yielded a nonsignificant result (*χ^2^* = 5.43, *p* = .14) and the AIC value for model *d2* (237.56) was lower than for *d1*. The residuals of models *d1* and *d2* were normally distributed, according to a visual inspection. Bonferroni-corrected pairwise comparisons showed significant differences between LE and the other four conditions, and between IE3 and IE2, HE and IE1, and HE and IE2 (Table 2). Non-significant differences with large (*d* > .8) effect sizes were found for IE3 and IE1. Non-significant differences with small effect sizes (0 < *d* < .02) were observed for the contrasts IE2 - IE1 and HE - IE3 (Figure 7a; parameters reported in supplementary file 2).

**Table 2.** Bonferroni-corrected pairwise contrast for *d’*-scores

| contrast | estimate | CI 2.5% | CI 97.5% | t | p | Cohen's d |
| --- | --- | --- | --- | --- | --- | --- |
| LE - IE1 | 0.55 | 0.07 | 1.02 | 3.16 | 0.02 | 0.98 |
| LE - IE2 | 0.54 | 0.07 | 1.01 | 3.11 | 0.02 | 0.96 |
| LE - IE3 | 1.04 | 0.56 | 1.51 | 5.99 | < 0.001 | 1.86 |
| LE - HE | 1.04 | 0.57 | 1.52 | 6.04 | < 0.001 | 1.86 |
| IE1 - IE2 | -0.01 | -0.48 | 0.46 | -0.04 | 1 | -0.02 |
| IE1 - IE3 | 0.49 | 0.02 | 0.96 | 2.84 | 0.05 | 0.88 |
| IE1 - HE | 0.5 | 0.03 | 0.97 | 2.88 | 0.04 | 0.89 |
| IE2 - IE3 | 0.5 | 0.03 | 0.97 | 2.88 | 0.04 | 0.89 |
| IE2 - HE | 0.51 | 0.03 | 0.98 | 2.93 | 0.03 | 0.91 |
| IE3 - HE | 0.01 | -0.46 | 0.48 | 0.05 | 1 | 0.02 |

**Figure 6.**
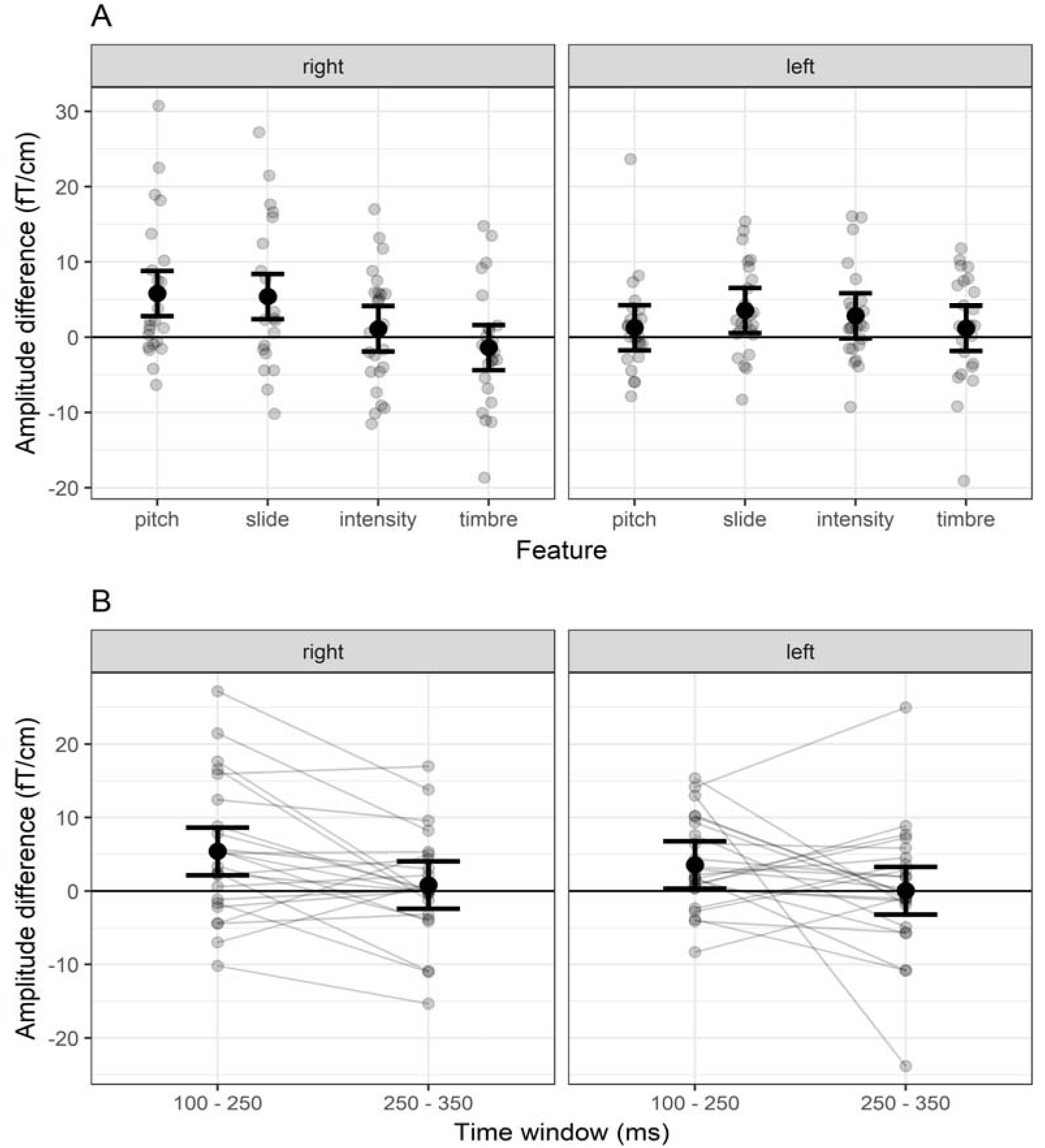
MMNm amplitude differences between low-entropy and high entropy conditions for (A) all features and (B) slide in two time windows, in both hemispheres. Error bars represent 95% confidence intervals.

Regarding *c*-scores, an intercept-only model (*cr0*) revealed a positive bias (figure 7b; supplementary file 2), suggesting an overall tendency for participants to answer negatively—i.e., no deviant present. A model with entropy as a categorical factor (*cr1*) explained the data significantly better (*χ^2^* = 11.74, *p* = .02) and had lower AIC (207.84) than the *cr0* (211.59) model. Pairwise comparisons revealed a significant difference only for the IE1 - IE3 contrast, although a large effect (*d* = .77) was also seen for the LE – IE3 contrast (Table 3).

**Table 3.** Bonferroni-corrected pairwise contrasts for *c*-scores

| contrast | estimate | CI 2.5% | CI 97.5% | t | p | Cohen's d |
| --- | --- | --- | --- | --- | --- | --- |
| LE - IE1 | -0.14 | -0.57 | 0.3 | -0.87 | 1 | -0.27 |
| LE - IE2 | 0.12 | -0.31 | 0.56 | 0.79 | 1 | 0.23 |
| LE - IE3 | 0.4 | -0.03 | 0.83 | 2.52 | 0.12 | 0.78 |
| LE - HE | 0.15 | -0.29 | 0.58 | 0.93 | 1 | 0.29 |
| IE1 - IE2 | 0.26 | -0.17 | 0.7 | 1.65 | 0.98 | 0.51 |
| IE1 - IE3 | 0.54 | 0.1 | 0.97 | 3.38 | 0.01 | 1.05 |
| IE1 - HE | 0.29 | -0.15 | 0.72 | 1.8 | 0.72 | 0.56 |
| IE2 - IE3 | 0.27 | -0.16 | 0.71 | 1.73 | 0.84 | 0.52 |
| IE2 - HE | 0.02 | -0.41 | 0.46 | 0.15 | 1 | 0.04 |
| IE3 - HE | -0.25 | -0.68 | 0.18 | -1.58 | 1 | -0.49 |

Regarding confidence ratings, the *c1* model (with a term for entropy) explained the data better (*χ^2^* = 236.88, *p* < .001) and had lower AIC (6816.9) than the *c0* (intercept-only) model (7045.8) (Figure 7c; supplementary file 2). The *c1* model also performed significantly better (*χ^2^* = 88.33, *p* < .001) and had lower AIC than the *c2* model (6899.3), which included entropy as an ordered variable. Adding random slopes improved model performance, as revealed by the comparison between *c1s* and *c1* (*χ^2^* = 234.02, *p* < .001), and *c2s* and *c2* (*χ^2^* = 101.36, *p* < .001). A comparison between c1s and *c2s* (*χ^2^* = 221, *p* < .001) showed that the best model was *c1s*—i.e. a model with a categorical effect of entropy and random-slopes. AIC values for *c1s* and *c2s* were 6610.9 and 6801.9, respectively. Pairwise comparisons for the *c1s* model revealed significant differences between LE and the other conditions (Table 4). No other comparisons were significant and yielded odds ratios smaller than 3 and larger than 0.5. In contrast, all pairwise comparisons for the *c1* model were significant, except between IE2 and HE, and IE1 and HE (Table 5). This indicates that differences between IE1, IE2, IE3 and HE are not detectable when individual variability in the relation between conditions is taken into account.

**Table 4.** Bonferroni-corrected pairwise contrasts for confidence ratings with no random slopes (model *c1*)

| contrast | odds ratio | CI 2.5% | CI 97.5% | z | p |
| --- | --- | --- | --- | --- | --- |
| LE - IE1 | 2.88 | 2.01 | 4.13 | 8.23 | < 0.001 |
| LE - IE2 | 4.27 | 2.97 | 6.14 | 11.21 | < 0.001 |
| LE - IE3 | 6.41 | 4.44 | 9.25 | 14.2 | < 0.001 |
| LE - HE | 4.05 | 2.84 | 5.79 | 11 | < 0.001 |
| IE1 - IE2 | 1.48 | 1.05 | 2.1 | 3.18 | 0.01 |
| IE1 - IE3 | 2.23 | 1.57 | 3.15 | 6.44 | < 0.001 |
| IE1 - HE | 1.41 | 1 | 1.98 | 2.81 | 0.05 |
| IE2 - IE3 | 1.5 | 1.06 | 2.12 | 3.29 | 0.01 |
| IE2 - HE | 0.95 | 0.67 | 1.34 | -0.43 | 1 |
| IE3 - HE | 0.63 | 0.45 | 0.89 | -3.76 | < 0.001 |

**Table 5.** Bonferroni-corrected pairwise contrasts for confidence ratings with random slopes (model *c1s*).

| contrast | odds ratio | CI 2.5% | CI 97.5% | z | p |
| --- | --- | --- | --- | --- | --- |
| LE - IE1 | 3.06 | 1.81 | 5.19 | 5.96 | < 0.001 |
| LE - IE2 | 5.34 | 2.12 | 13.45 | 5.09 | < 0.001 |
| LE - IE3 | 8.33 | 2.63 | 26.37 | 5.16 | < 0.001 |
| LE - HE | 4.93 | 2.23 | 10.9 | 5.65 | < 0.001 |
| IE1 - IE2 | 1.74 | 0.63 | 4.81 | 1.54 | 1 |
| IE1 - IE3 | 2.72 | 0.77 | 9.59 | 2.23 | 0.26 |
| IE1 - HE | 1.61 | 0.65 | 4.02 | 1.46 | 1 |
| IE2 - IE3 | 1.56 | 0.94 | 2.58 | 2.48 | 0.13 |
| IE2 - HE | 0.92 | 0.62 | 1.37 | -0.56 | 1 |
| IE3 - HE | 0.59 | 0.34 | 1.03 | -2.65 | 0.08 |

**Figure 7.**
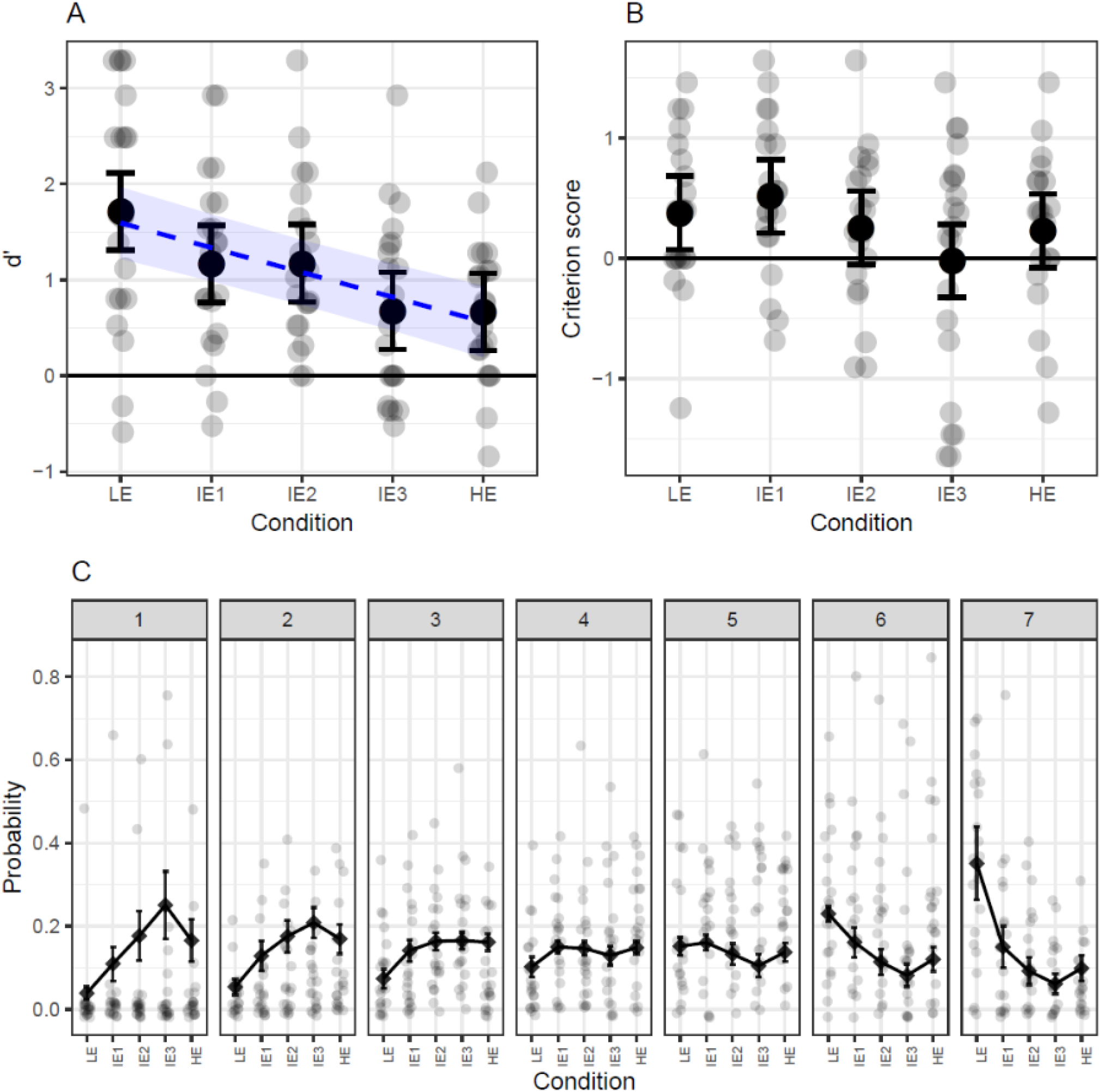
Plots of A) *d’*-scores, B) criterion scores and C) confidence ratings with their respective parameter estimates and 95% confidence intervals. For *d*’-scores the slope from the *d2* model (with entropy as an ordered linear predictor) is also plotted as a blue dashed line. Confidence ratings are presented as the probability of responses in each category (1-7) as a function of the different entropy conditions. LE = low entropy, IE = intermediate entropy, HE = high entropy.

## 4. Discussion

In the present study, we investigated whether prediction error responses are affected by uncertainty in auditory contexts that are more complex, ecologically valid and real-sounding than those typically used in neuroimaging research. Employing tone sequences that resembled real music, we found decreased MMNm amplitudes for pitch and slide deviants in high-entropy (HE) as compared to low-entropy (LE) stimuli. This modulation was paralleled by accuracy scores—and, to some extent, confidence ratings—in a behavioral deviance detection task, which tended to decrease with higher entropy levels. These findings are in agreement with theories of predictive processing (Clark, 2013; Feldman & Friston, 2010; Hohwy, 2012) and models of musical expectations (Hansen et al., 2017; Ross & Hansen, 2016; Vuust et al., 2018) which propose that prediction error responses are reduced in contexts with low as compared to high predictive precision.

Our results are consistent with empirical research already showing reduced auditory prediction error responses in uncertain contexts (Garrido et al., 2013; Sohoglu & Chait, 2016; Southwell & Chait, 2018). They are also in agreement with studies that found differences in sustained tonic activity—as opposed to phasic responses such as the MMN—when comparing low- and high-entropy contexts (Auksztulewicz et al., 2017; Barascud et al., 2016; Nastase, Iacovella, & Hasson, 2014; Overath et al., 2007). Closer to assessing the effect of precision in a musical context, recent research shows that the entropy of short rhythmic sequences modulates MMN responses (Lumaca, Haumann, Brattico, Grube, & Vuust, 2019). However, in this study rhythms were presented as repeated short patterns, which makes them less akin to actual musical stimuli than our HE sequences. Together with studies showing an absence of MMN responses in random contexts (Hsu et al., 2015; Jacobsen & Schröger, 2001; Maess et al., 2007), the evidence suggests a role of precision in auditory processing at the neural level. However, our study is the first to suggest that precision might also modulate prediction error during music listening—a common, highly structured, and more ecologically valid auditory context.

### 4.1. Sources of uncertainty

In our experiments, differences in uncertainty arise from two main sources. First, the degree of repetitiveness of the stimuli. For example, in LE sequences, a simple pattern is iterated, which allows very precise expectations about upcoming events. In contrast, the scarcity of exact repetitions in HE sequences makes it harder to predict specific continuations from the preceding tones. This can be seen in the IDyOM estimates: for LE, IC drops after the first occurrence of the pattern, whereas HE sequences tend to have higher IC levels throughout (supplementary file 1). This is also the case for the intermediate conditions in which repetitiveness decreased, going from an iterative pattern to completely random sequences. The fact that we found differences in *d’*-scores between IE1 and IE3—where the pitch alphabet was the same—suggests that repetitiveness itself is sufficient to affect deviance detection.

The other source of uncertainty is pitch alphabet. For example, in the LE condition, where the alphabet consisted of three pitch heights, the probability of each tone was higher on average than in IE1 sequences, where the pitch alphabet consisted of eight pitch heights. Therefore, the larger the alphabet, the higher the uncertainty of the context. The fact that we found differences in *d’*-scores between LE and IE1—which are equally repetitive but have different alphabets—indicates that this factor is also sufficient to modulate the detection of unexpected sounds. Importantly, in the MEG experiment the difference in pitch alphabet was minimized by transposing LE sequences to cover the same range as HE sequences. Therefore, it is tempting to conclude that differences in neural activity are not driven by pitch alphabet but rather by repetitiveness. However, it is possible that participants learned the local alphabet of each 32-note sequence instead of the global alphabet of all the sequences. This would be reinforced by the transpositions, assuming that participants heard the sequences with respect to the tonal center of the respective key. Further research could aim to disentangle the contributions of repetitiveness and pitch alphabet to the perceived context uncertainty and its effect on prediction error at the neural level.

### 4.2. IDyOM estimates

IDyOM’s quantitative estimates generally followed our qualitative manipulations. Entropy values were lower for LE than for the IE conditions, and in turn these were lower than for HE. However, some nonlinearities can be observed. Specifically, LE seemed to yield much lower values than the other four conditions, whereas differences between IE1, IE2, IE3 and HE were more subtle. Considering that we have relatively complex and realistic stimuli and that IDyOM was not used to make quantitative predictions, these nonlinearities do not affect the overall conclusions of the study. Instead, they help us understand how well the model captures context uncertainty in more realistic settings. IC also tended to follow the expected pattern, with higher values for contexts with higher uncertainty. The only exception is the HE condition, which had lower values than all the intermediate conditions. This might reflect the fact that small pitch intervals—i.e. 1 or 2 semitones—which have high probability in most Western tonal music, were more common in HE than the IE conditions. Therefore, these results constitute an interesting example of how entropy and information content can sometimes be dissociated. In sum, IDyOM’s estimates seem to characterize reasonably well the uncertainty of a context (cf. Hansen & Pearce, 2014). Note, however, that the estimates are based on an adjustment which was meant to account for the differences in pitch alphabet between conditions. Without the adjustment, the estimations are unreliable and lead to misleading results such as LE sequences having higher entropy than IE3 (random) sequences (supplementary figure 3). Nonetheless, we regard the adjustment as the best approximation to the simulation of a listener who learns the alphabet of a context, taking into account that such learning has not been implemented in IDyOM yet.

Since IDyOM was used here descriptively as a way to characterize the stimuli, the insight we can gain from it is limited by several factors. First, we used low-level deviant sounds which currently are not accounted for by the model and do not happen in real music. We acknowledge that directly modeling uncertainty, prediction error and their interaction for the case of unexpected tones that actually occur in music might provide a better insight on their neural and behavioral manifestations. However, the deviants employed here have the advantage that they allow a clear dissociation between prediction error and uncertainty, since they are equally unexpected regardless of the context, as they are always outside the reference distribution. A related issue is that, even though IDyOM modeled pitch predictions, our pitch deviants—i.e. out-of-tune tones—were not included in the model, which might lead some to argue that IDyOM’s estimates and the pitch MMNm address predictive processes of different kinds. A possible mechanism linking these seemingly distinct levels of processing could be categorical perception (Goldstone & Hendrickson, 2010; Schulze, 1989; Siegel & Siegel, 1977), in which mistuned tones would be heard as renditions of the closest pitch category in the tuning system, and therefore be affected by its uncertainty. Moreover, an interesting future path of enquiry would be to use single-tone entropy and IC estimates to assess the effect of uncertainty on prediction error through regression and single-trial analyses. Such an approach was not used here, as our aims were different. We were interested in the uncertainty of the context as a whole, rather than its moment by moment fluctuations. We also tried to maximize the contrast between LE and HE stimuli to observe clear differences in neural activity. Furthermore, we wanted to assess the behavior of the MMN in a more complex and real-sounding setting, which is why the comparison with an existing MMN paradigm was a natural choice. Therefore, our work can be seen as a step towards more detailed models of the effect of precision on prediction error.

### 4.3. Feature-specific effects

The observed MMNm amplitudes seemed to be more affected by entropy for pitch and slide than for timbre and intensity deviants. We attribute this to the fact that we manipulated entropy in the pitch dimension, while other dimensions were restricted to the same two standard and deviant values (piano timbre vs. telephone receiver; high intensity vs. low intensity) across all conditions. The fact that these differences were most prominent in right temporal gradiometers might be due to the MMNm signal being largest in these sites, which in turn is consistent with the rightward asymmetry for music processing (e.g. Brattico et al., 2006; Koelsch et al., 2000; Zatorre, Belin, & Penhune, 2002). Consequently, it seems that stimulus uncertainty particularly affected deviants that depend on pitch information, which points to a feature-specific effect of precision on the MMN. This interpretation is consistent with MMN recordings in multi-feature (Näätänen, Pakarinen, Rinne, & Takegata, 2004; Vuust et al., 2011) and no-standard (Kliuchko, Heinonen-Guzejev, Vuust, Tervaniemi, & Brattico, 2016; Pakarinen et al., 2010) paradigms in which auditory regularities are created for a specific feature even though sounds constantly change in other features.

Furthermore, the suggested feature-selectivity is particularly interesting in the case of the slide MMNm, which had an unusual shape that extended and peaked beyond 250 ms. This shape may be attributed to the fact that, unlike in previous experiments, the pitch glide spanned the whole duration of the tone and thus the MMNm amplitude seemed to mirror the increasing magnitude of the continuous deviation. The fact that only the earlier portion of the response was different between conditions might reflect the coexistence of two violations. Since the slide deviant started two semitones below its corresponding standard, we propose that the first section is a pitch MMNm, while the second corresponds to a proper slide MMNm. Thus, in the LE block, where there were much more precise pitch expectations, slide deviants were heard first as a “wrong” pitch and afterwards as a pitch glide. In contrast, for HE sequences, the sense of a “wrong” pitch would be weaker but the glide would be equally surprising. If this account is correct, the fact that the first (pitch) but not the second (slide proper) part is reduced for HE is consistent with the idea of a feature-specific precision modulation of the MMNm. In any case, the differences between features discussed above have to be taken with caution since they constitute a non-hypothesized finding. Future work manipulating uncertainty across different features and measuring its effects on different types of deviants is required to properly test the proposed feature-selectivity.

### 4.4. Behavioral experiment

In the behavioral experiment, both *d’*-scores and confidence ratings were lower for HE than LE sequences. This confirms the MEG results and suggests that the effect of precision on neural prediction error for pitch-related features is associated with a reduced ability to distinguish pitch deviants from regular sounds. Criterion scores showed that participants were generally biased towards not identifying the deviants, and that there were no big differences between conditions except for IE3, which was less biased than IE1 and seemingly LE. This might suggest that when participants are faced with a completely random context, they are less conservative and guess more. However, given the absence of a consistent pattern for other conditions, this difference has to be interpreted with caution.

Analyses of *d’*-scores revealed that even fine-grained differences in context uncertainty can affect deviance detection. This is the case of comparisons such as LE-IE1 or IE2-IE3. As discussed above, these contrasts also show that both repetitiveness and pitch alphabet are sufficient for the effect to be elicited. In general, conditions with higher entropy tended to yield lower *d’*-scores. This is further supported by the *d2* model, in which the five entropy conditions were included as an ordered linear predictor. Since this model has similar likelihood—as suggested by the non-significant likelihood ratio test—, but also less parameters and a lower AIC value than the *d*1 (categorical) model, the results support the idea that higher uncertainty leads to reduced deviance detection even for small manipulations of the context. However, it has to be noted that, for the categorical model, virtually no differences were found between IE1 and IE2, and IE3 and HE, which slightly departs from a decreasing linear trend and IDyOM’s estimates. The reason for this pattern are not easily identifiable in the current design, but it is possible that stimulus-specific variation played a role, since conditions had a different number of unique sequences or melodies.

Regarding confidence scores, clear differences were found between LE and the other four conditions. In other words, participants tended to give higher ratings in the context with the highest precision. This suggests that both neural and accuracy measures are related to a subjective feeling of certainty. Interestingly, differences among the other conditions were observable in the case of the *c1* model, which did not include random slopes, but not the *c1s* model, which included them. Thus, the apparent differences between subtle changes in context uncertainty seemed to be driven only by a few subjects and disappear when individual differences are taken into account. This indicates that subjective certainty is less sensitive to contextual factors than deviance detection itself. It is worth noting that both *d’*-scores and confidence showed large differences between LE and the other conditions, but either smaller or no differences between IE1, IE2, IE3 and HE. This goes in line with the nonlinearities in IDyOM estimates, and suggests that precision is maximized in repetitive contexts with small pitch alphabets.

Taken together, MEG and behavioral results point to a precision modulation of prediction error. However, the relationship between the two experiments has to be taken with caution since the behavioral task required participants to actively detect deviations, whereas in the MEG session they listened to the sounds passively while watching a silent movie. Thus, there were additional higher-order processes involved in the former which means that differences in *d’*-scores and confidence ratings cannot be ascribed exclusively to the processes reflected in the MMN. Further research involving active tasks and neurophysiological recordings is needed to assess the contribution of different components and processing stages to the effect of interest.

### 4.5. Limitations and future directions

The work presented here has some limitations. For example, in the MEG experiment, we compared two types of stimuli that differ in several aspects. As mentioned before, both repetitiveness and pitch-alphabet seemed to play a role, and even though they influence the entropy of the sequences, it is not possible to properly disentangle their individual contributions here. Another aspect is the repetition rate of individual sequences. In the LE condition of the MEG experiment, individual sequences were repeated every two trials, whereas individual melodies in the HE condition were repeated every twelve trials. This could have created a stronger long-term memory representation for LE sequences which might be responsible at least for part of the effect. Something similar might have happened in the behavioral experiment. However, this possible confound cannot be regarded as the only explanation for our results since we found differences for conditions with equal repetition rates (e.g. LE and IE1). This interpretation is also compatible with the precision-based explanation if one regards long-term representations as very precise expectations. Moreover, even though acoustic confounds were minimized in the MEG experiment by using the same pitch alphabet in both conditions, it is still possible that differences in pitch distributions between conditions created acoustic differences, as pitch discrimination is more difficult at very low or very high frequency ranges (Sek & Moore, 1995). However, this explanation is unlikely since in both conditions the range of frequencies was similarly covered (supplementary figure 1) and pitches were not higher than 5 kHz, which is the frequency range for which discrimination significantly decreases.

Another aspect to consider is to what extent the findings can be generalized outside the conditions of the experiment. For example, even though our stimuli are much more real-sounding than in most research, they are still far from actual music. This comes from the use of isochronous sounds, the lack of artistic expression, the somewhat excessive repetition, the absence of concurrent sound streams, and the introduction of deviants that do not occur in real music. Nonetheless, we regard the stimuli as realistic enough to be considered musical, and as a good compromise between experimental control and ecological validity. A related issue is that our participants watched a movie, which arguably is not the most common setting for music listening. Thus, how uncertainty affects prediction error under attentive or other listening conditions remains to be seen. Another caveat is that participants were all nonmusicians and were recruited from an online database, which might have introduced bias in the sample. Therefore, replications with different populations are needed. Finally, we acknowledge that our hypotheses and methods were not pre-registered, which is not desirable if one aims to minimize analytical bias. However, in the spirit of transparency, we have shared our data and code openly so that our work can be directly reproduced, scrutinized and built upon by the research community.

Despite these shortcomings, we believe our study makes two main contributions to the literature. First, our HE condition constitutes an advance in the multi-feature experimental paradigms used to study prediction error responses through the MMN. With it, we demonstrated that it is possible to obtain reliable—albeit reduced—MMN signals with more complex and realistic stimuli in a multi-feature paradigm. We have done so by exploiting the possibilities of abstract-feature MMN responses (Paavilainen, 2013), which arise, not from the violation of an exact sensory representation established through the exact repetition of a stimulus, but from the breach of an abstract regularity established in a constantly changing acoustic stream.

Second and most importantly, our results show how prediction error signals behave in more complex and more ecologically valid auditory contexts than those typically studied. This has consequences for the current knowledge and future directions of research in audition and music cognition. For example, some musical styles, such as atonal music, exploit uncertainty as an artistic resource. Therefore, one could hypothesize imprecise predictions and reduced prediction error in these styles, something that could be studied with the new MMN paradigm reported here. This is relevant for the understanding of the aesthetic and emotional experiences associated with these types of music (Mencke, Omigie, Wald-Fuhrmann, & Brattico, 2019). Another point of interest is how individual factors play a role in predictive processing in uncertain contexts. For instance, it has been suggested that musical expertise enhances the precision of auditory predictive models (e.g., Vuust et al., 2018). Thus, one could hypothesize that the effect of precision on neural prediction error would be less pronounced in musical experts and potentially be modulated by stylistic expertise. Finally, the fact that the MMN seems to be reduced and in some cases disappears in uncertain contexts might question whether this brain response fully represents the core processes involved in auditory prediction. Arguably, expectations are still generated in complex settings, which means that other neural processes might also be at play. In that sense, the use of more ecologically valid stimuli in combination with computational models such as IDyOM could reveal more accurately the neural mechanisms involved.

## 5. Conclusion

In this study, we investigated whether prediction error responses are modulated by uncertainty in more complex and real-sounding auditory contexts than those typically studied. Our results show that prediction error responses—as indexed by the MMNm, accuracy scores and, to some extent, confidence ratings—are reduced in contexts with higher uncertainty and suggest that this reduction may be constrained to features that depend on the auditory dimension whose uncertainty is manipulated (pitch in our case). Thus, in line with recent theories of predictive processing, our work provides further support to precision-weighted prediction error as a fundamental principle for brain function, and moves us a step closer to understanding auditory predictive processing in the rich environments of daily life.

## Design and Analysis Transparency (21-word solution)

We report how we determined our sample size, all data exclusions (if any), all data inclusion/exclusion criteria, whether inclusion/exclusion criteria were established prior to data analysis, all manipulations, and all measures in the study.

## Acknowledgments

We wish to thank the project initiation group, namely Christopher Bailey, Torben Lund and Dora Grauballe, for their help with setting up the experiments. We also thank Nader Sedghi, Massimo Lumaca, Giulia Donati, Ulrika Varankaité, Giulio Carraturo, Riccardo Proietti, and Claudia Iorio for assistance during MEG recordings. We are indebted as well to the group of Italian trainees from IISS Simone-Morea, Conversano, who helped with the behavioral experiment. Finally, we thank Hella Kastbjerg for checking the English language of this manuscript. The Center for Music in the Brain is funded by the Danish National Research Foundation (DNRF 117), which did not have any influence on the scientific content of this article.

Declaration of interests: none

