## Supplementary File 2 for "Reduced prediction error responses in high- as compared to low-uncertainty musical contexts"

### Summary of linear mixed models

In this supplemantary file, we report all the models and parameters refered to in the article.

1. Model of MGA differences by feature:

$$MGA = \beta_0 + \beta_{feature} + \beta_{hemisphere} + \beta_{feature:hemisphere} + \sigma_{subject} + \sigma_{residual}$$

| parameter | estimate | 2.5 % | 97.5 % |
|---|---|---|---|
| Intercept | 5.80 | 2.79 | 8.81 |
| feature: slide | -0.40 | -3.96 | 3.17 |
| feature: intensity | -4.68 | -8.24 | -1.11 |
| feature: timbre | -7.18 | -10.75 | -3.62 |
| hemisphere: left | -4.55 | -8.12 | -0.99 |
| feat*hem: slide/left | 2.71 | -2.34 | 7.75 |
| feat*hem: intensity/left | 6.26 | 1.22 | 11.31 |
| feat*hem: timbre/left | 7.13 | 2.08 | 12.17 |
| SD (subject) | 4.03 | 2.76 | 5.88 |
| SD (residual) | 6.27 | 5.65 | 7.00 |

2. Model of differences in slide between conditions

$$MGA_{slide} = \beta_0 + \beta_{latency} + \beta_{hemisphere} + \beta_{latency:hemisphere} + \sigma_{subject} + \sigma$$

| parameter | estimate | 2.5 % | 97.5 % |
|---|---|---|---|
| Intercept | 5.40 | 2.24 | 8.57 |
| latency: 250-350 | -4.58 | -8.58 | -0.58 |
| hemisphere: left | -1.85 | -5.85 | 2.15 |
| lat*hem: 250-350/left | 1.08 | -4.58 | 6.74 |
| SD (subject) | 3.60 | 1.11 | 5.83 |
| SD (residual) | 7.13 | 5.98 | 8.30 |

3. Models of d'-scores

$$d_0 = \beta_0 + \sigma_{subject} + \sigma$$

| parameter | estimate | 2.5 % | 97.5 % |
|---|---|---|---|
| Intercept | 1.08 | 0.73 | 1.43 |
| SD (subject) | 0.72 | 0.50 | 1.05 |
| SD (residual) | 0.71 | 0.61 | 0.83 |

$$d_1 = \beta_0 + \beta_{entropy} + \sigma_{subject} + \sigma$$

| parameter | estimate | 2.5 % | 97.5 % |
|---|---|---|---|
| Intercept | 1.71 | 1.30 | 2.12 |
| entropy: IE1 | -0.55 | -0.89 | -0.20 |
| entropy: IE2 | -0.54 | -0.88 | -0.20 |
| entropy: IE3 | -1.04 | -1.38 | -0.69 |
| entropy: HE | -1.04 | -1.39 | -0.70 |
| SD (subject) | 0.74 | 0.54 | 1.07 |
| SD (residual) | 0.56 | 0.48 | 0.66 |

$$d_2 = \beta_0 + \beta * entropy + \sigma_{subject} + \sigma$$

| parameter | estimate | 2.5 % | 97.5 % |
|---|---|---|---|
| Intercept | 1.85 | 1.43 | 2.27 |
| entropy | -0.26 | -0.34 | -0.18 |
| SD (subject) | 0.74 | 0.54 | 1.07 |
| SD (residual) | 0.58 | 0.50 | 0.68 |

4. Models of criterion scores:

$$cr_0 = \beta_0 + \sigma_{subject} + \sigma$$

| parameter | estimate | 2.5 % | 97.5 % |
|---|---|---|---|
| Intercept | 0.27 | 0.03 | 0.51 |
| SD (subject) | 0.48 | 0.32 | 0.71 |
| SD (residual) | 0.55 | 0.48 | 0.65 |

$$cr_1 = \beta_0 + \beta_{entropy} + \sigma_{subject} + \sigma$$

| parameter | estimate | 2.5 % | 97.5 % |
|---|---|---|---|
| Intercept | 0.38 | 0.07 | 0.69 |
| entropy: IE1 | 0.14 | -0.18 | 0.45 |
| entropy: IE2 | -0.12 | -0.44 | 0.19 |
| entropy: IE3 | -0.40 | -0.71 | -0.08 |
| entropy: HE | -0.15 | -0.46 | 0.17 |
| SD (subject) | 0.48 | 0.33 | 0.71 |
| SD (residual) | 0.51 | 0.45 | 0.60 |

5. Cumulative link models (ordinal regression)

$$c_0 : \log \frac{p(confidence)}{1 - p(confidence)} = \theta_{confidence} - \sigma_{subject} - \sigma$$

| parameter | odds | 2.5 % | 97.5 % |
|---|---|---|---|
| 1\|2 | 0.02 | 0.01 | 0.04 |
| 2\|3 | 0.09 | 0.05 | 0.16 |
| 3\|4 | 0.30 | 0.17 | 0.52 |
| 4\|5 | 0.74 | 0.42 | 1.31 |
| 5\|6 | 2.23 | 1.27 | 3.90 |
| 6\|7 | 8.76 | 4.95 | 15.49 |
| SD (subject) | 3.67 | NA | NA |

$$c_1 : \log \frac{p(confidence)}{1 - p(confidence)}_0 = \theta_{confidence} - \beta * entropy - \sigma_{subject} - \sigma$$

| parameter | odds | 2.5 % | 97.5 % |
|---|---|---|---|
| 1\|2 | 0.01 | 0.00 | 0.01 |
| 2\|3 | 0.02 | 0.01 | 0.05 |
| 3\|4 | 0.08 | 0.04 | 0.16 |
| 4\|5 | 0.22 | 0.12 | 0.41 |
| 5\|6 | 0.73 | 0.39 | 1.35 |
| 6\|7 | 3.19 | 1.72 | 5.99 |
| IE1 | 0.35 | 0.27 | 0.44 |
| IE2 | 0.23 | 0.18 | 0.30 |
| IE3 | 0.16 | 0.12 | 0.20 |
| HE | 0.25 | 0.19 | 0.32 |
| SD (subject) | 3.97 | NA | NA |

$$c_2 : \log \frac{p(confidence)}{1 - p(confidence)} = \theta_{confidence} - \beta * entropy - \sigma_{subject} - \sigma$$

| parameter | odds | 2.5 % | 97.5 % |
|---|---|---|---|
| 1\|2 | 0.01 | 0.00 | 0.01 |
| 2\|3 | 0.03 | 0.02 | 0.05 |
| 3\|4 | 0.10 | 0.05 | 0.18 |
| 4\|5 | 0.26 | 0.14 | 0.47 |
| 5\|6 | 0.80 | 0.44 | 1.48 |
| 6\|7 | 3.42 | 1.86 | 6.23 |
| entropy | 0.71 | 0.67 | 0.75 |
| SD (subject) | 3.82 | NA | NA |

$$c_{1s} : \log \frac{p(confidence)}{1 - p(confidence)} = \theta_{confidence} - \beta_{entropy} - \sigma_{subject} - \sigma_{entropy*subject} - \sigma$$

| parameter | odds | 2.5 % | 97.5 % |
|---|---|---|---|
| 1\|2 | 0.00 | 0.00 | 0.01 |
| 2\|3 | 0.02 | 0.01 | 0.04 |
| 3\|4 | 0.06 | 0.03 | 0.14 |
| 4\|5 | 0.18 | 0.09 | 0.39 |
| 5\|6 | 0.66 | 0.31 | 1.40 |
| 6\|7 | 3.35 | 1.58 | 7.17 |
| IE1 | 0.33 | 0.23 | 0.47 |
| IE2 | 0.19 | 0.10 | 0.36 |
| IE3 | 0.12 | 0.05 | 0.27 |
| HE | 0.20 | 0.12 | 0.35 |
| SD (subject) | 5.47 | NA | NA |
| SD (IE1) | 1.86 | NA | NA |
| SD (IE2) | 3.97 | NA | NA |
| SD (IE3) | 5.93 | NA | NA |
| SD (HE) | 3.16 | NA | NA |

$$c_{2s} : \log \frac{p(confidence)}{1 - p(confidence)} = \theta_{confidence} - \beta * entropy - \sigma_{subject} - \sigma_{entropy*subject} - \sigma$$

| parameter | odds | 2.5 % | 97.5 % |
|---|---|---|---|
| 1\|2 | 0.00 | 0.00 | 0.01 |
| 2\|3 | 0.02 | 0.01 | 0.05 |
| 3\|4 | 0.08 | 0.04 | 0.18 |
| 4\|5 | 0.22 | 0.10 | 0.49 |
| 5\|6 | 0.73 | 0.33 | 1.60 |
| 6\|7 | 3.25 | 1.48 | 7.24 |
| entropy | 0.68 | 0.57 | 0.81 |
| SD (subject) | 6.05 | NA | NA |
| SD (entropy) | 1.48 | NA | NA |
