## Supplementary File 1 for "Reduced prediction error responses in high- as compared to low-uncertainty musical contexts"

Individual stimulus sequences used in the experiments with their respective IDyOM estimates

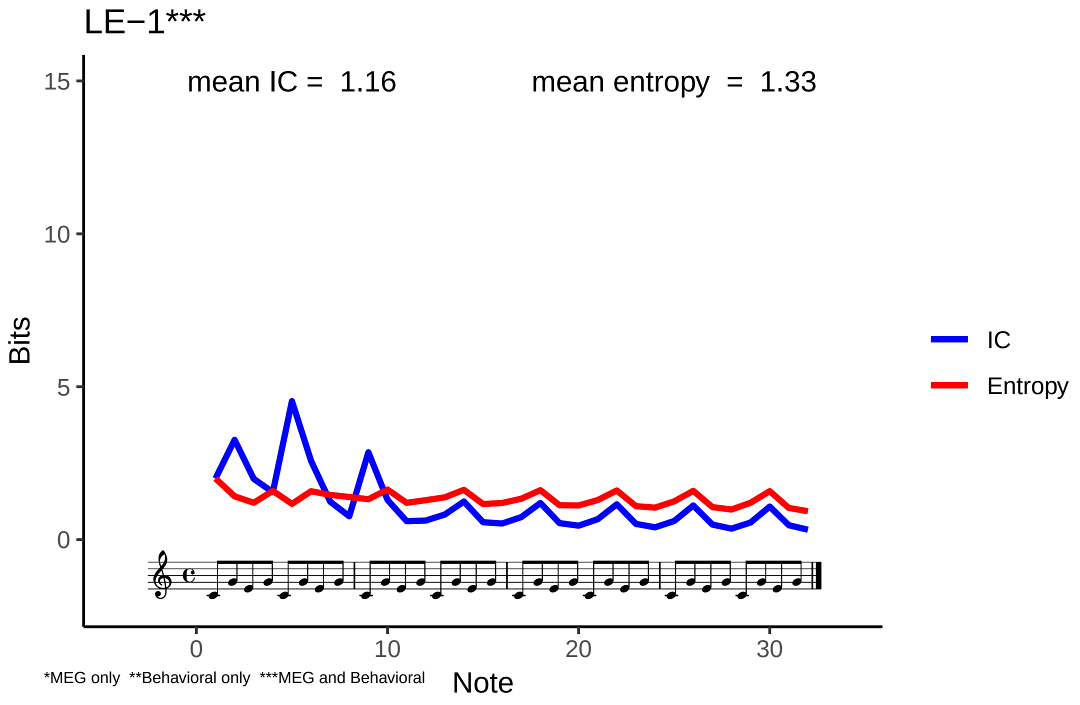

LE−1***
mean IC = 1.16
mean entropy = 1.33
Bits
15
10
5
0
0
10
20
30
Note
IC
Entropy
*MEG only **Behavioral only ***MEG and Behavioral

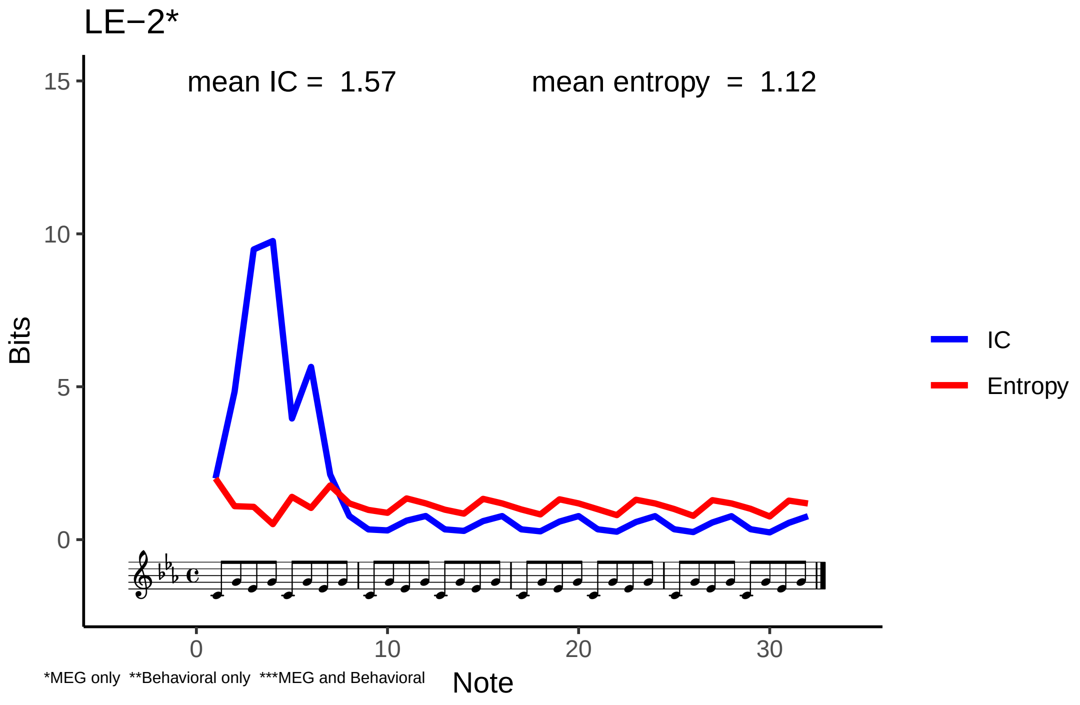

LE−2*
mean IC = 1.57
mean entropy = 1.12
Bits
15
10
5
0
0
10
20
30
Note
IC
Entropy
*MEG only **Behavioral only ***MEG and Behavioral

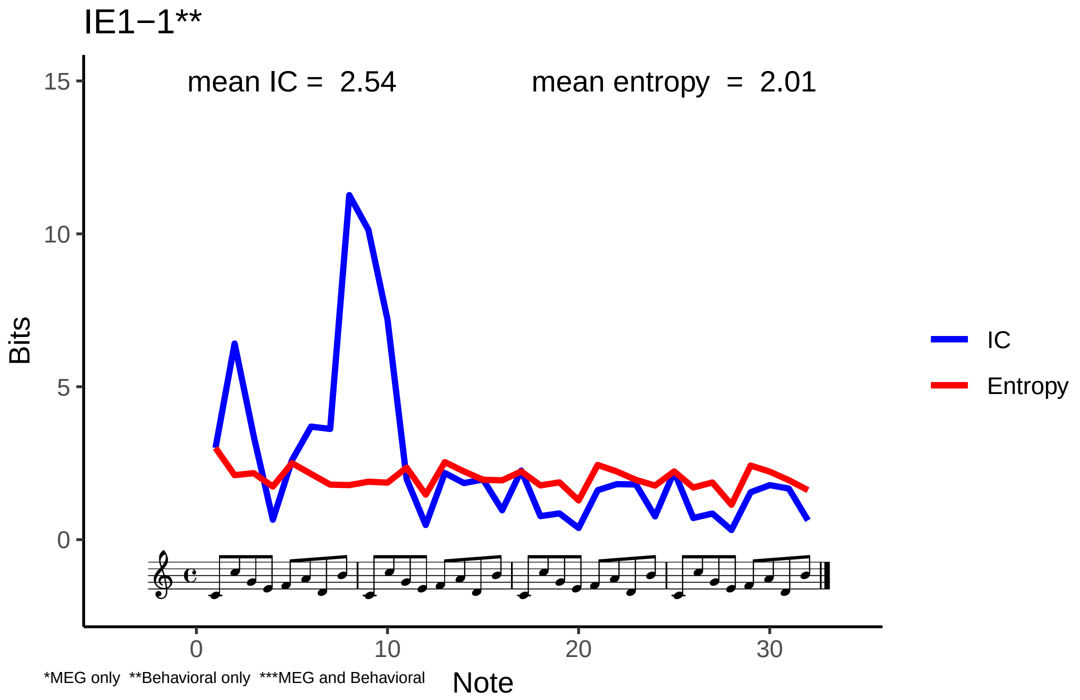

IE1−1**
mean IC = 2.54
mean entropy = 2.01
Bits
15
10
5
0
0
10
20
30
Note
IC
Entropy
*MEG only **Behavioral only ***MEG and Behavioral

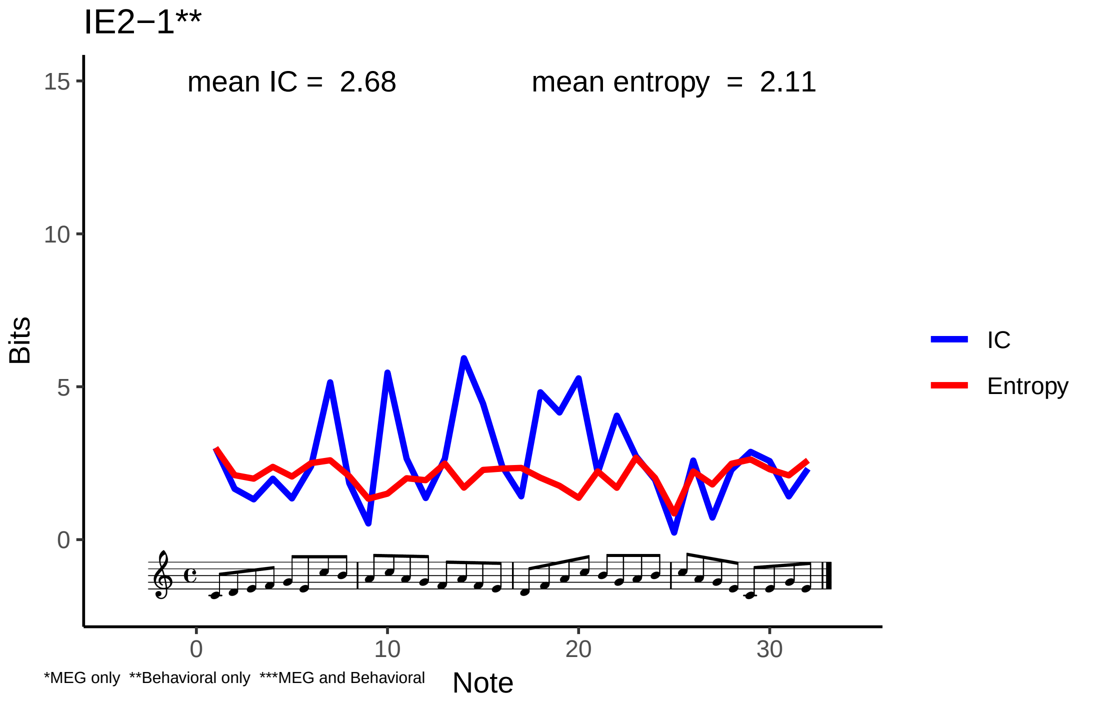

IE2−1**
mean IC = 2.68
mean entropy = 2.11
Bits
15
10
5
0
0
10
20
30
Note
IC
Entropy
*MEG only **Behavioral only ***MEG and Behavioral

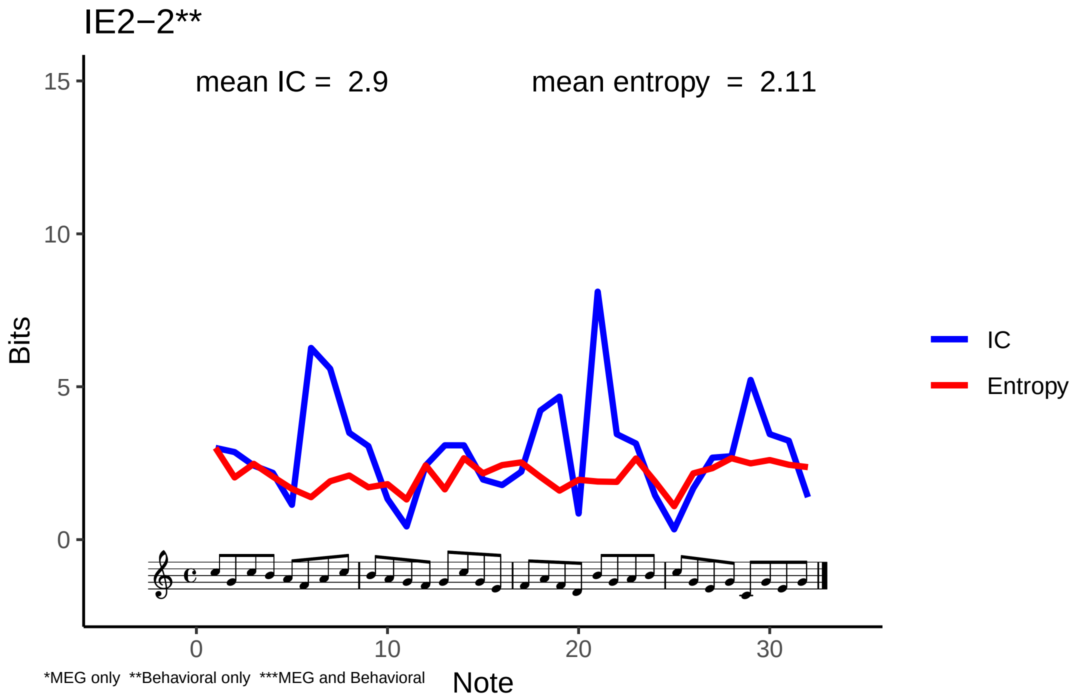

IE2−2**
mean IC = 2.9
mean entropy = 2.11
Bits
15
10
5
0
0
10
20
30
Note
IC
Entropy
*MEG only **Behavioral only ***MEG and Behavioral

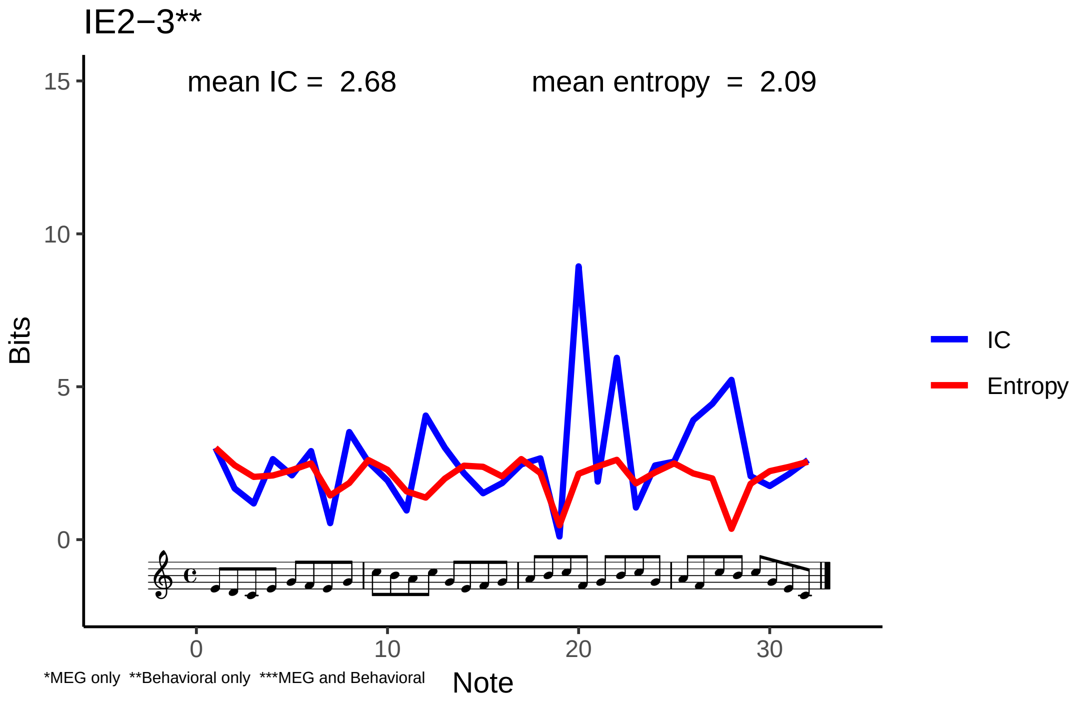
IE2−3**
mean IC = 2.68
mean entropy = 2.09
Bits
15
10
5
0
0
10
20
30
Note
IC
Entropy
*MEG only **Behavioral only ***MEG and Behavioral

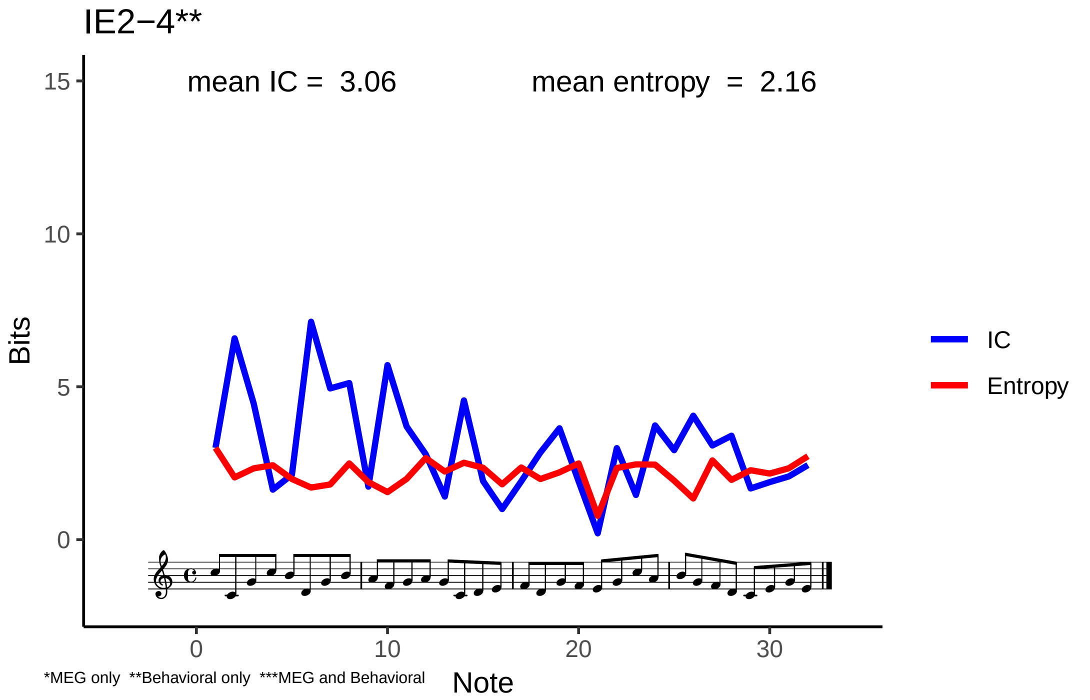

IE2−4**
mean IC = 3.06
mean entropy = 2.16
Bits
15
10
5
0
IC
Entropy
0
10
20
30
Note
*MEG only **Behavioral only ***MEG and Behavioral

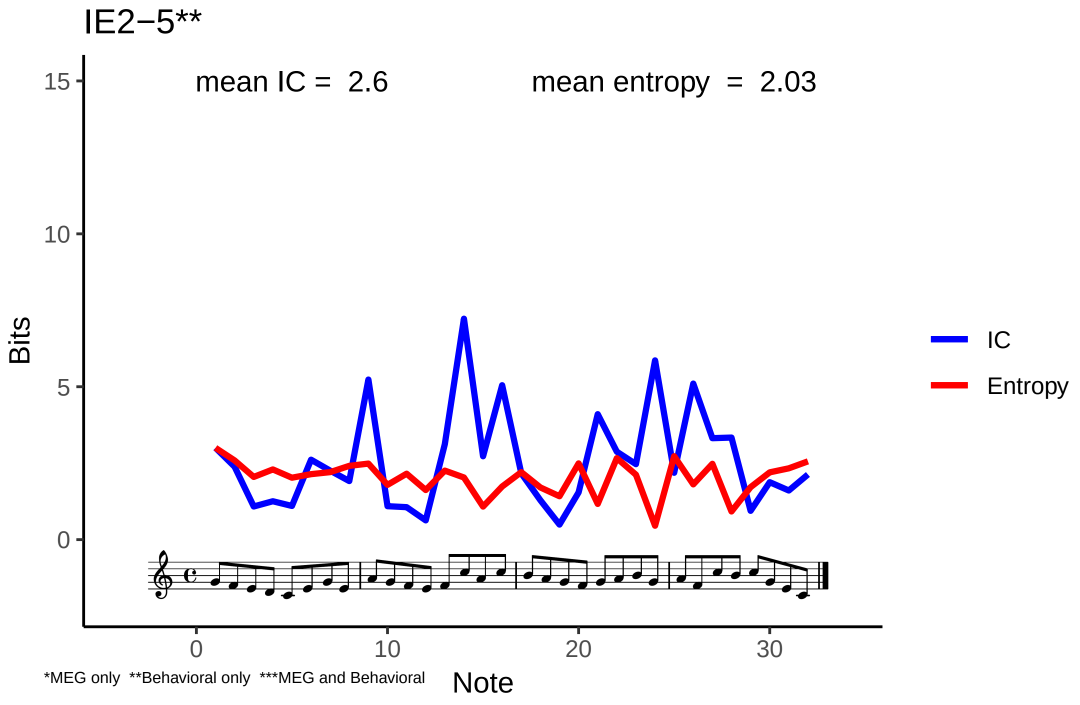
IE2−5**
mean IC = 2.6
mean entropy = 2.03
15
10
5
0
Bits
0
10
20
30
Note
IC
Entropy
*MEG only **Behavioral only ***MEG and Behavioral

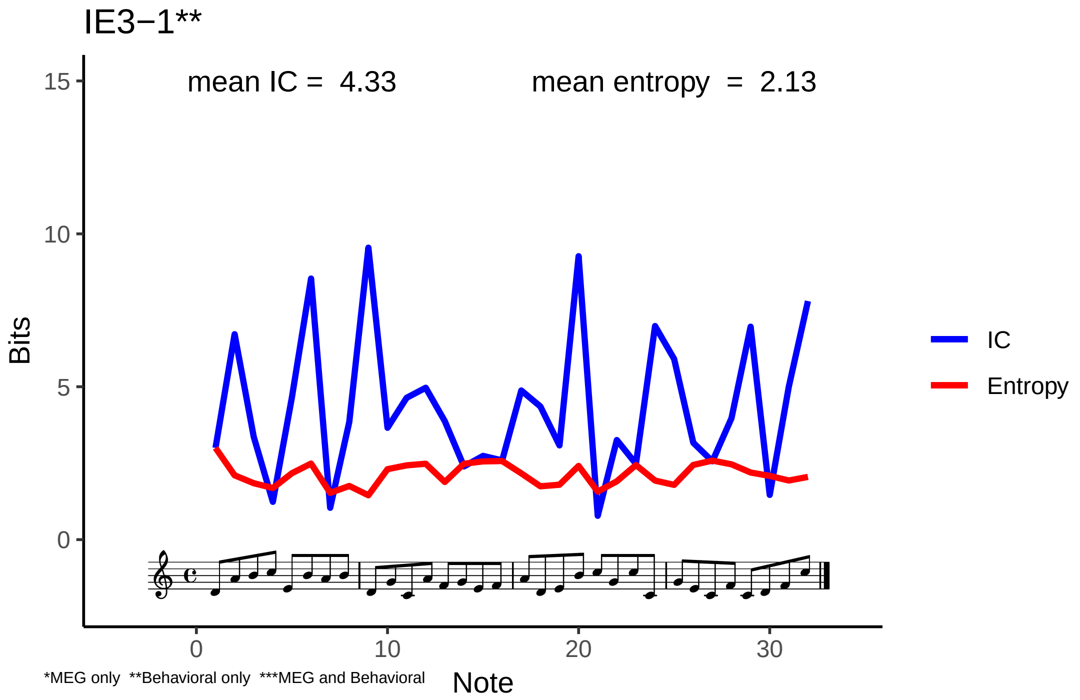

IE3−1**
mean IC = 4.33
mean entropy = 2.13
Bits
15
10
5
0
0
10
20
30
Note
IC
Entropy
*MEG only **Behavioral only ***MEG and Behavioral

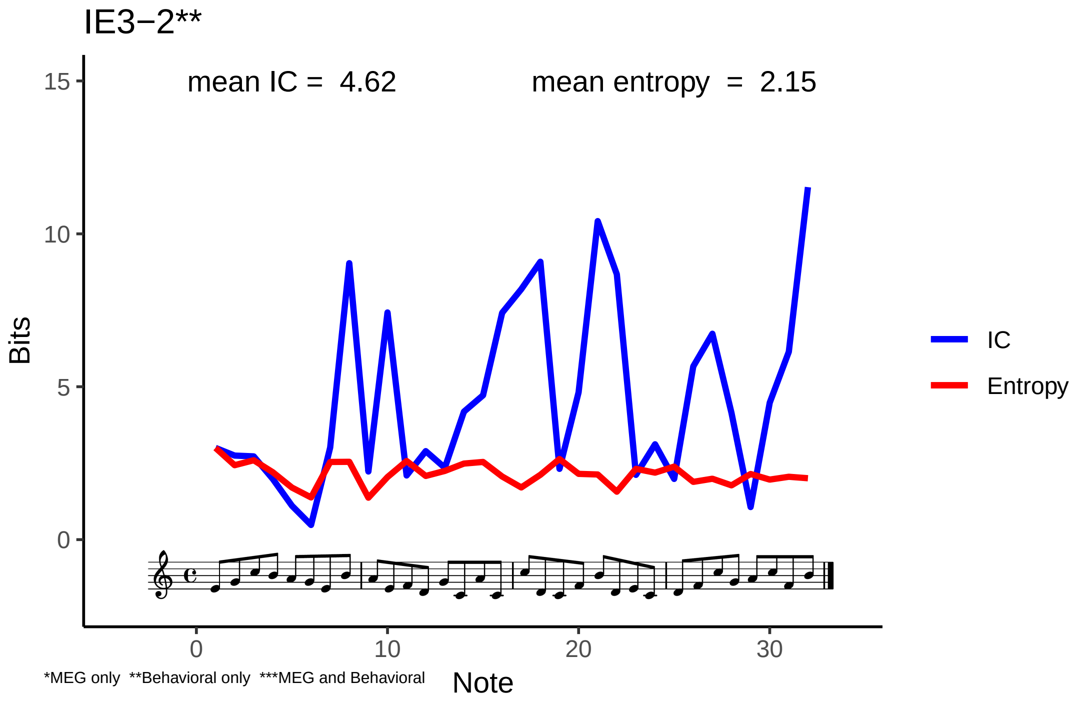

IE3−2**
mean IC = 4.62
mean entropy = 2.15
Bits
15
10
5
0
0
10
20
30
Note
IC
Entropy
*MEG only **Behavioral only ***MEG and Behavioral

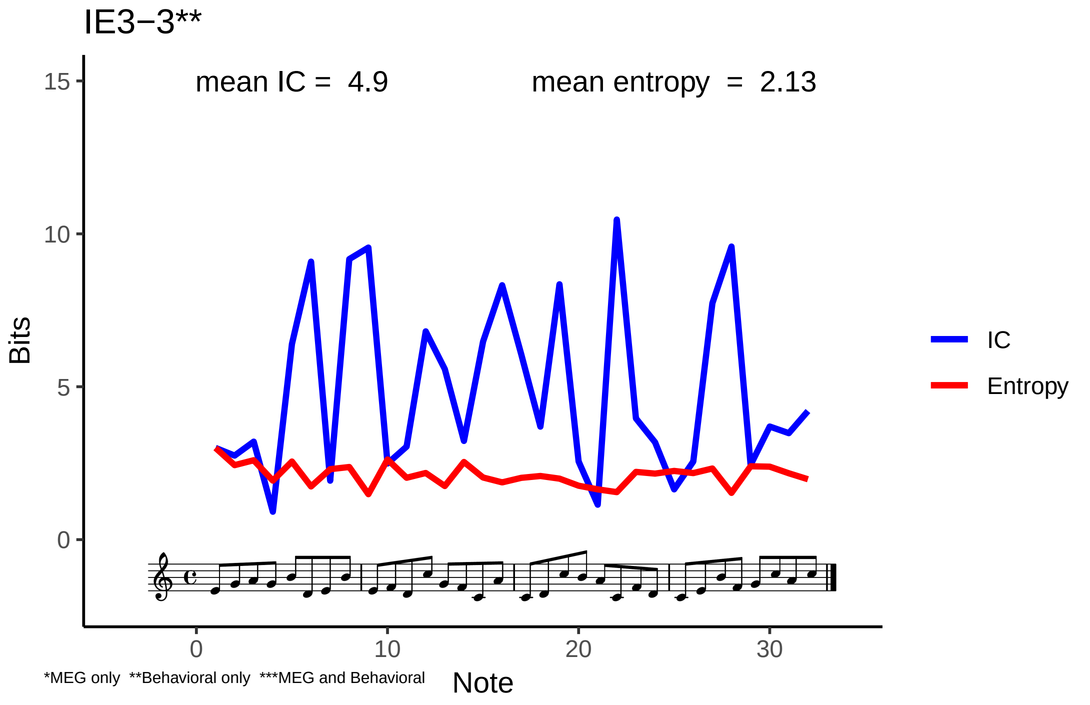

IE3−3**
mean IC = 4.9
mean entropy = 2.13
Bits
15
10
5
0
0
10
20
30
Note
IC
Entropy
*MEG only **Behavioral only ***MEG and Behavioral

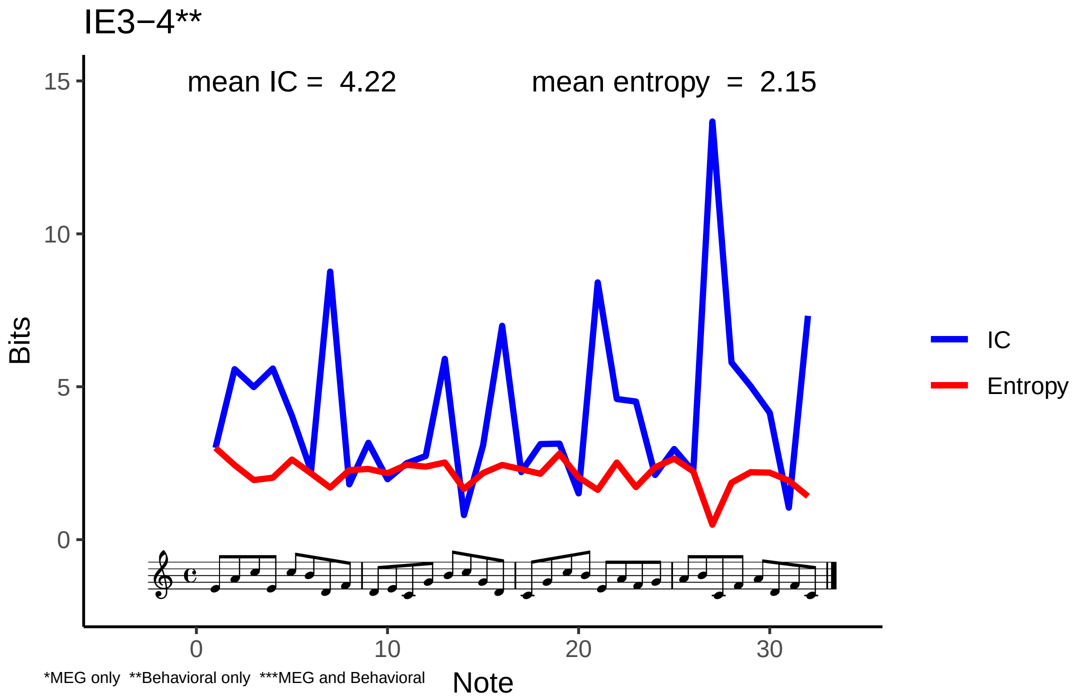
IE3−4**
mean IC = 4.22
mean entropy = 2.15
Bits
15
10
5
0
0
10
20
30
Note
IC
Entropy
*MEG only **Behavioral only ***MEG and Behavioral

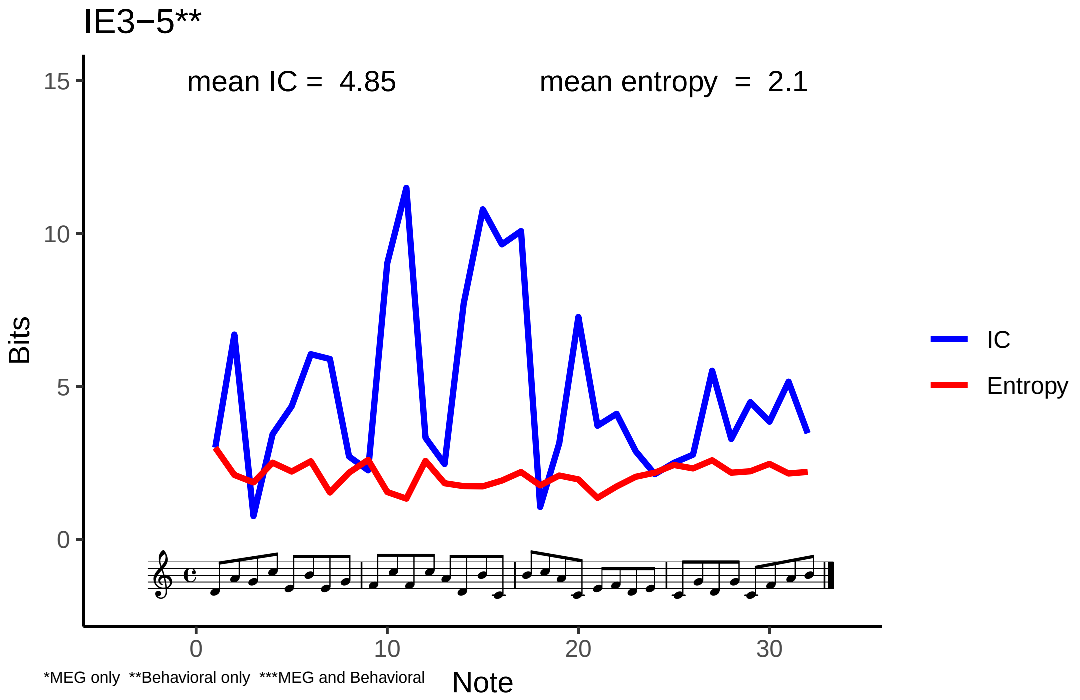

IE3−5**
mean IC = 4.85
mean entropy = 2.1
Bits
15
10
5
0
0
10
20
30
Note
IC
Entropy
*MEG only **Behavioral only ***MEG and Behavioral

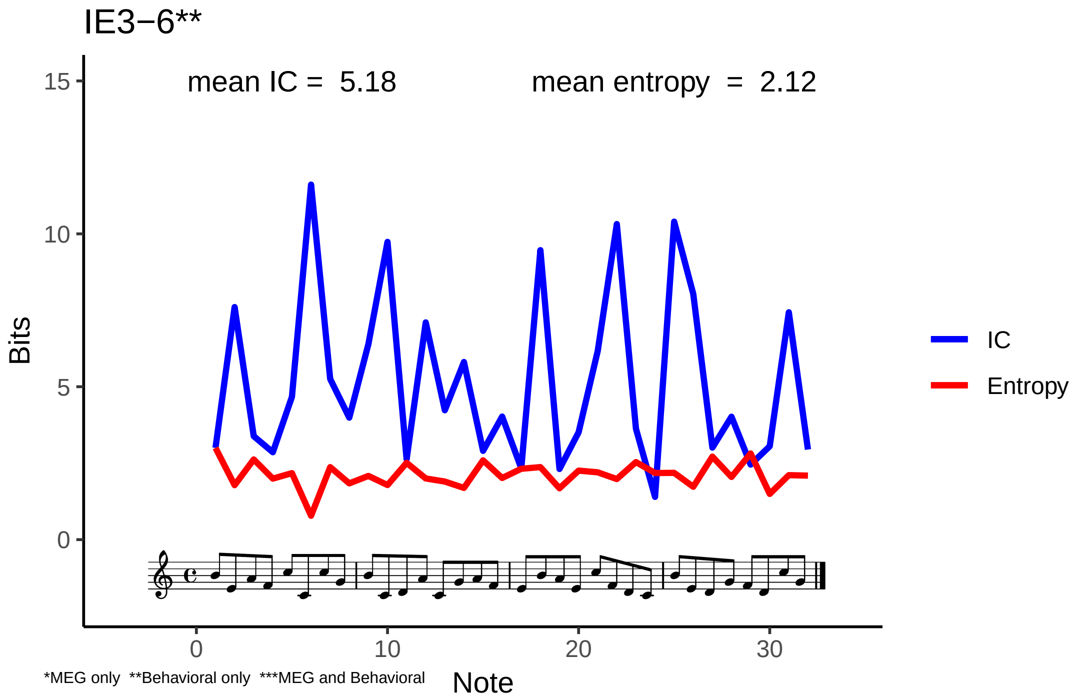

IE3−6**
mean IC = 5.18
mean entropy = 2.12
Bits
15
10
5
0
0
10
20
30
Note
IC
Entropy
*MEG only **Behavioral only ***MEG and Behavioral

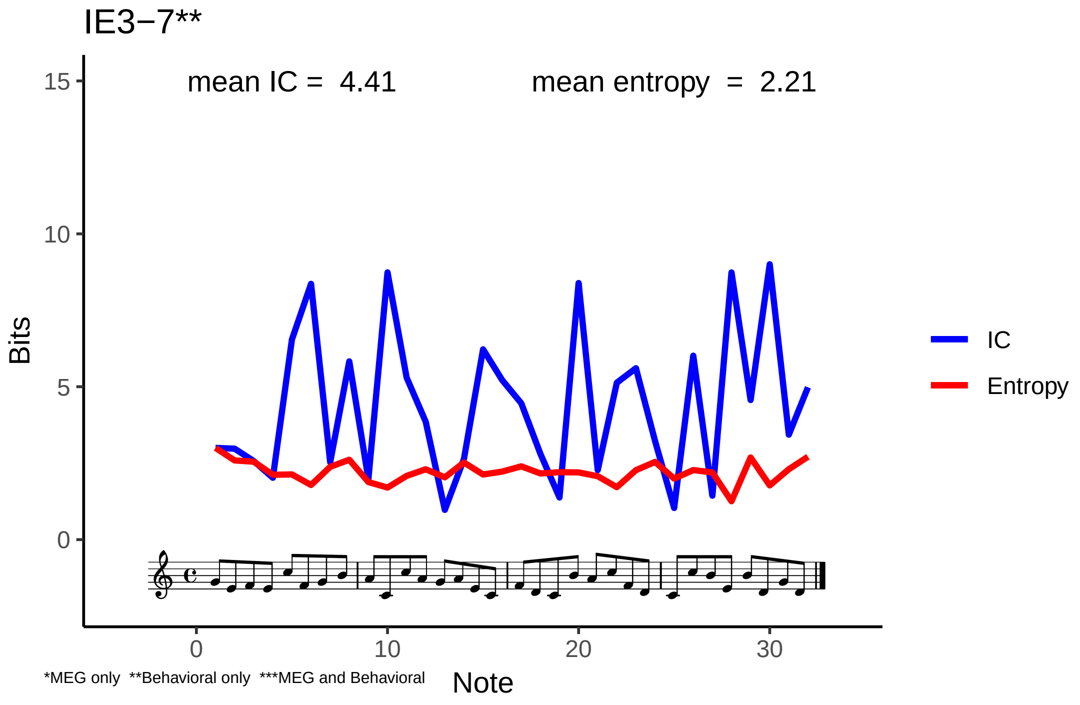

IE3−7**
mean IC = 4.41
mean entropy = 2.21
Bits
15
10
5
0
0
10
20
30
Note
IC
Entropy
*MEG only **Behavioral only ***MEG and Behavioral

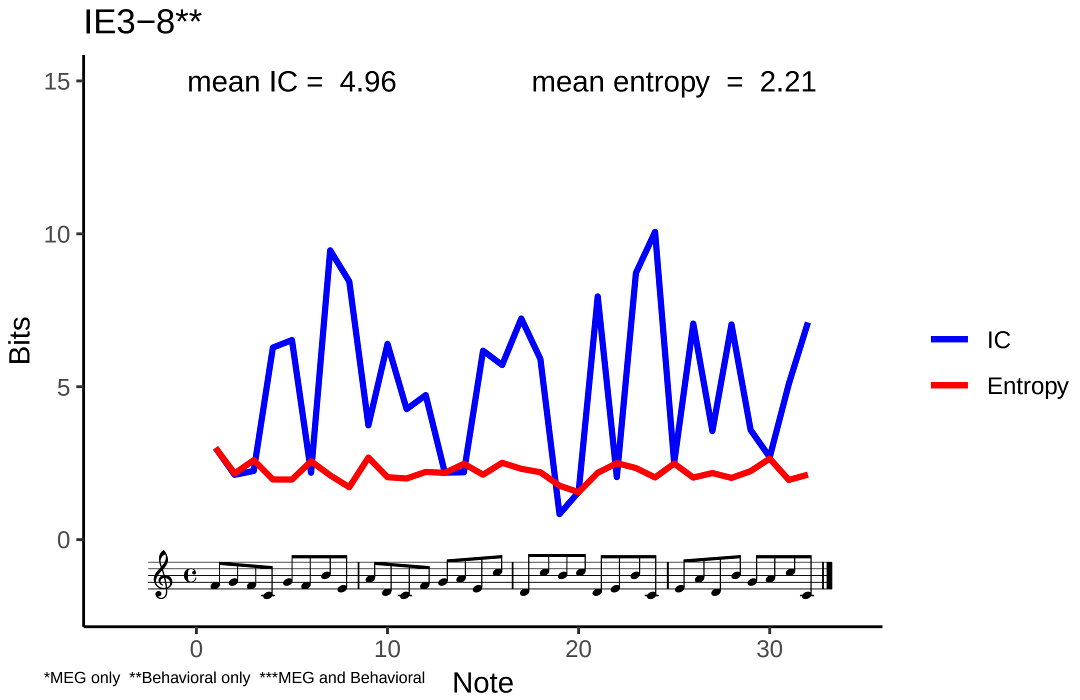
IE3−8**
mean IC = 4.96
mean entropy = 2.21
15
10
5
0
Bits
IC
Entropy
0
10
20
30
Note
*MEG only **Behavioral only ***MEG and Behavioral

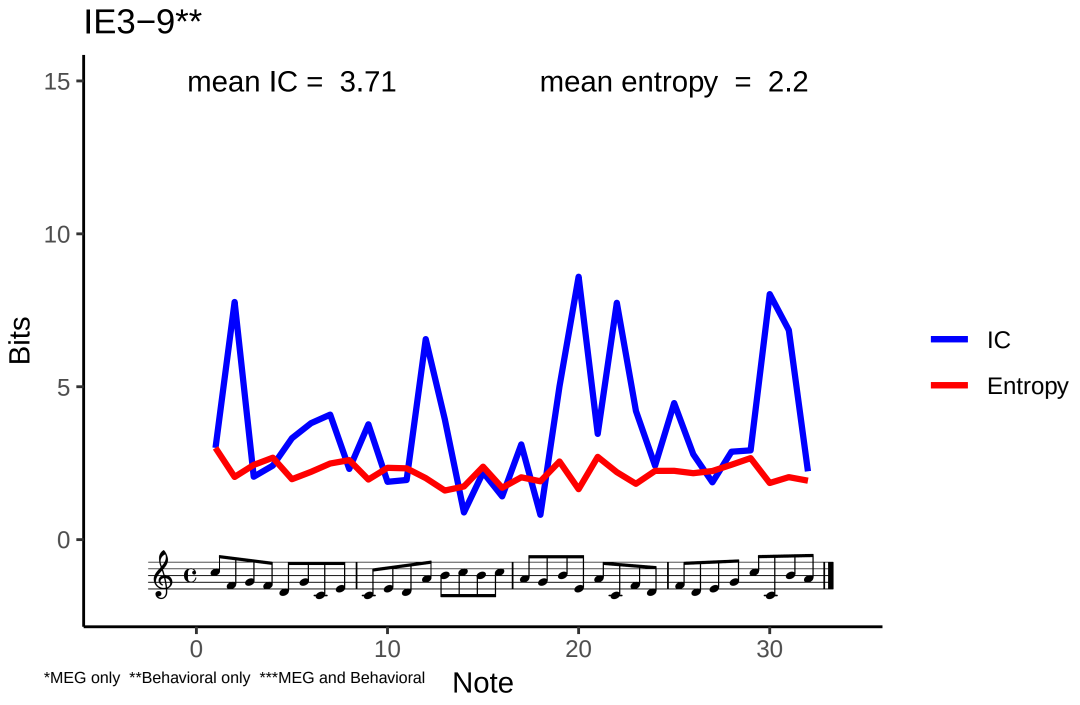

IE3−9**
mean IC = 3.71
mean entropy = 2.2
Bits
15
10
5
0
0
10
20
30
Note
IC
Entropy
*MEG only **Behavioral only ***MEG and Behavioral

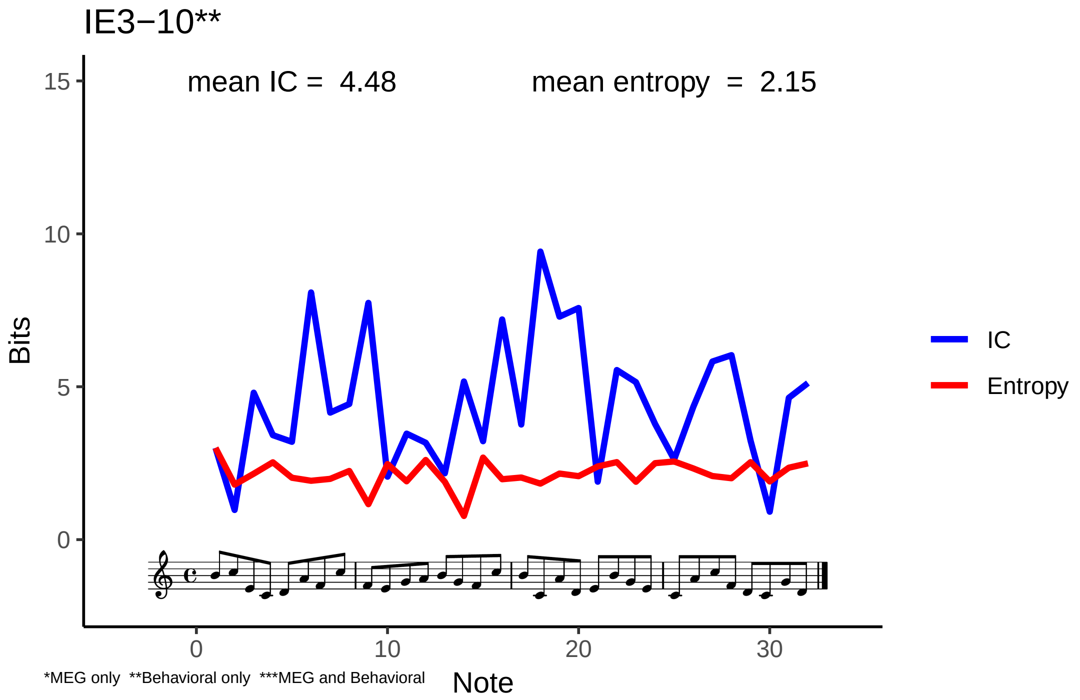
IE3−10**
mean IC = 4.48
mean entropy = 2.15
Bits
15
10
5
0
0
10
20
30
Note
IC
Entropy
*MEG only **Behavioral only ***MEG and Behavioral

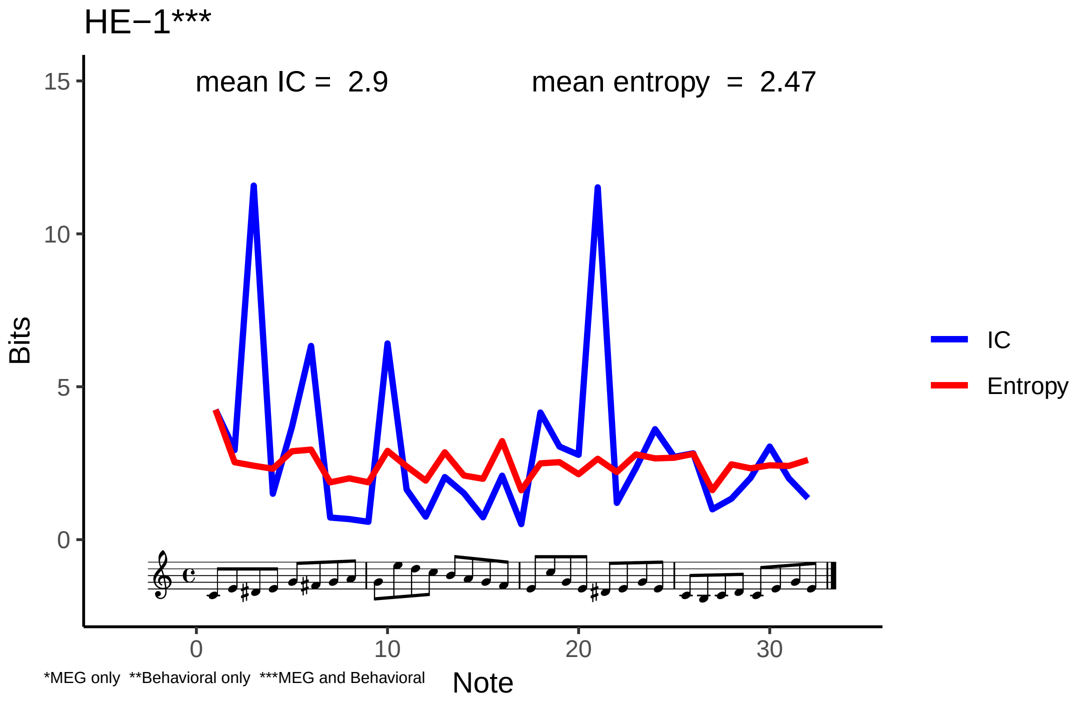
HE−1***
mean IC = 2.9
mean entropy = 2.47
Bits
15
10
5
0
0
10
20
30
Note
IC
Entropy
*MEG only **Behavioral only ***MEG and Behavioral

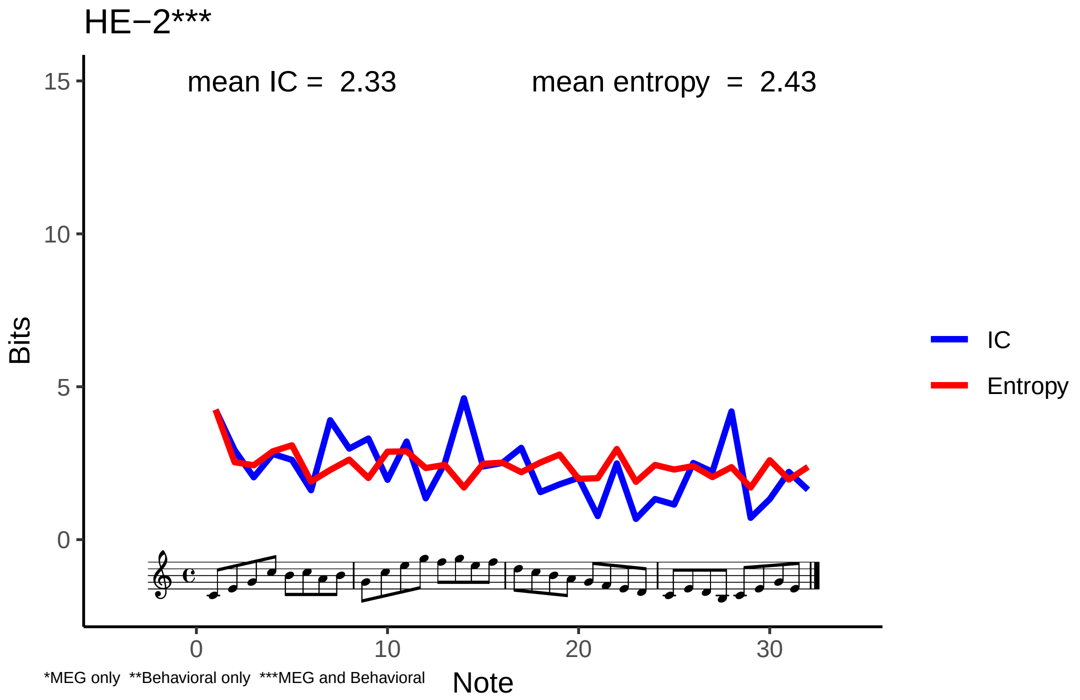
HE−2***
mean IC = 2.33
mean entropy = 2.43
Bits
15
10
5
0
0
10
20
30
Note
IC
Entropy
*MEG only **Behavioral only ***MEG and Behavioral

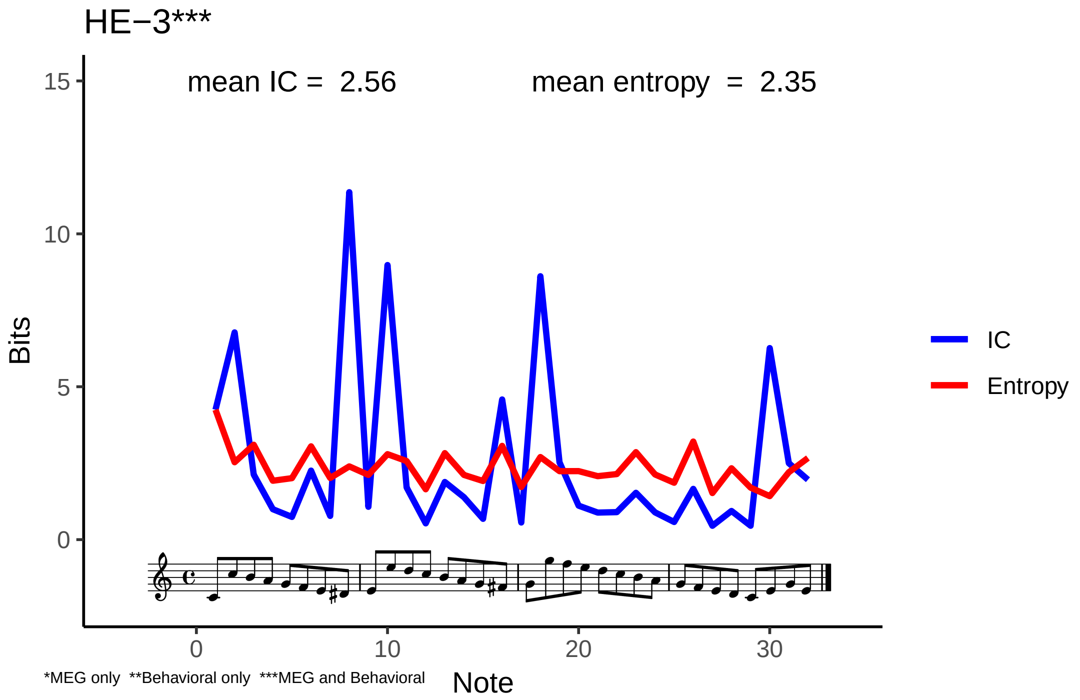

HE−3***
mean IC = 2.56
mean entropy = 2.35
Bits
15
10
5
0
0
10
20
30
Note
IC
Entropy
*MEG only **Behavioral only ***MEG and Behavioral

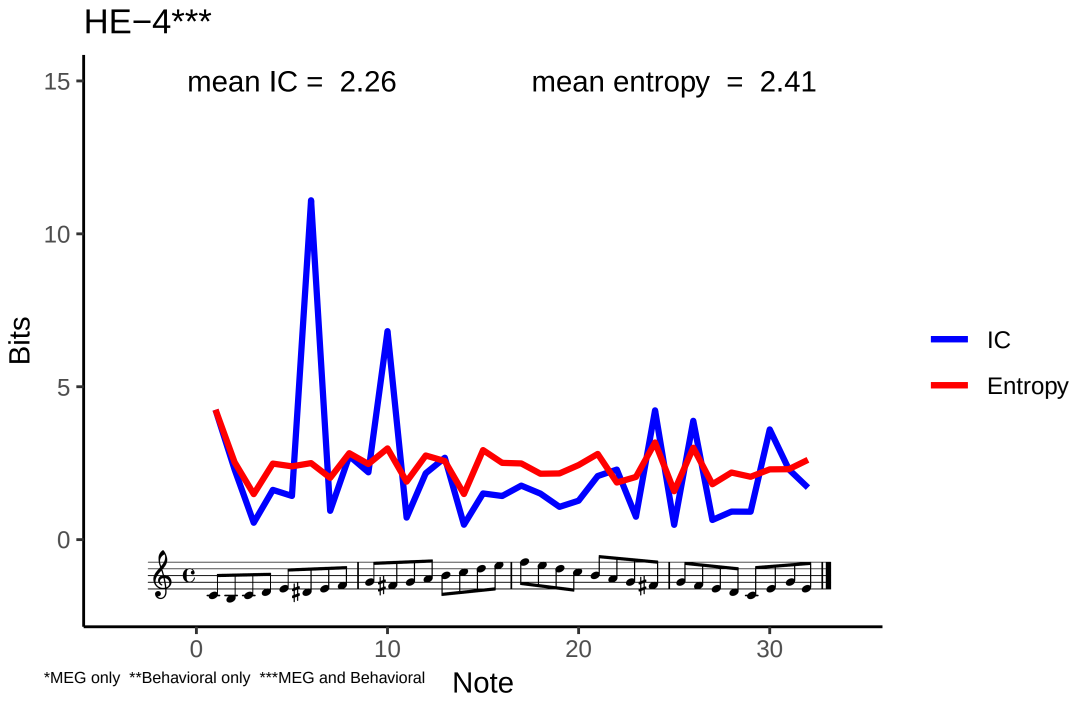
HE−4***
mean IC = 2.26
mean entropy = 2.41
Bits
15
10
5
0
0
10
20
30
Note
IC
Entropy
*MEG only **Behavioral only ***MEG and Behavioral

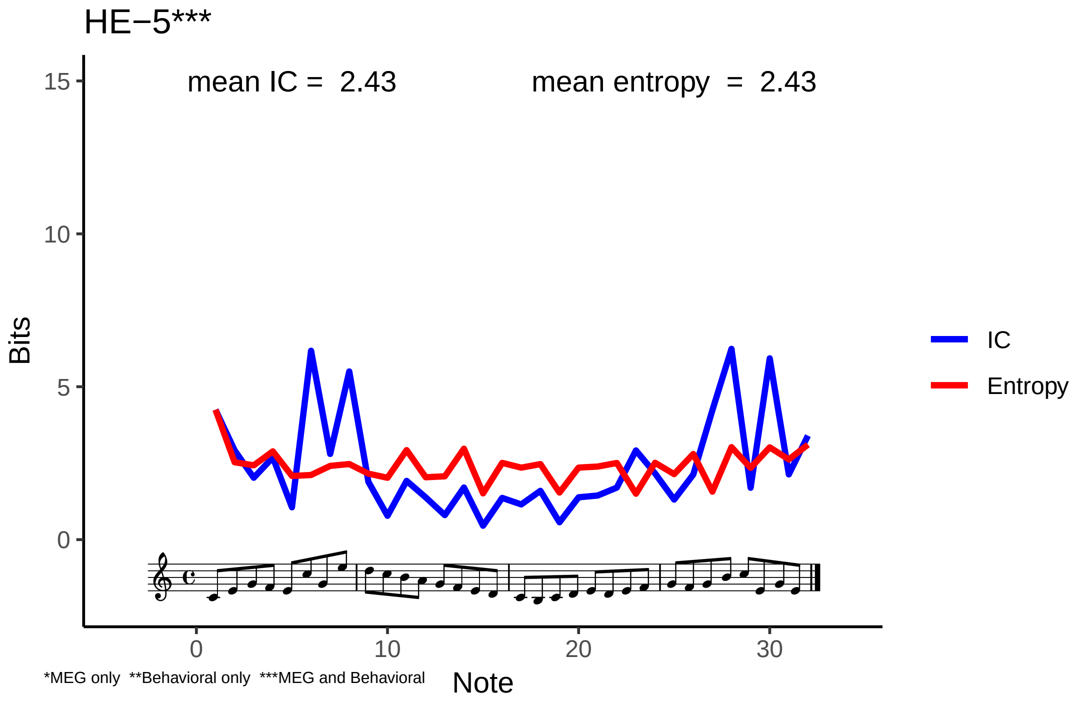
HE−5***
mean IC = 2.43
mean entropy = 2.43
Bits
15
10
5
0
0
10
20
30
Note
IC
Entropy
*MEG only **Behavioral only ***MEG and Behavioral

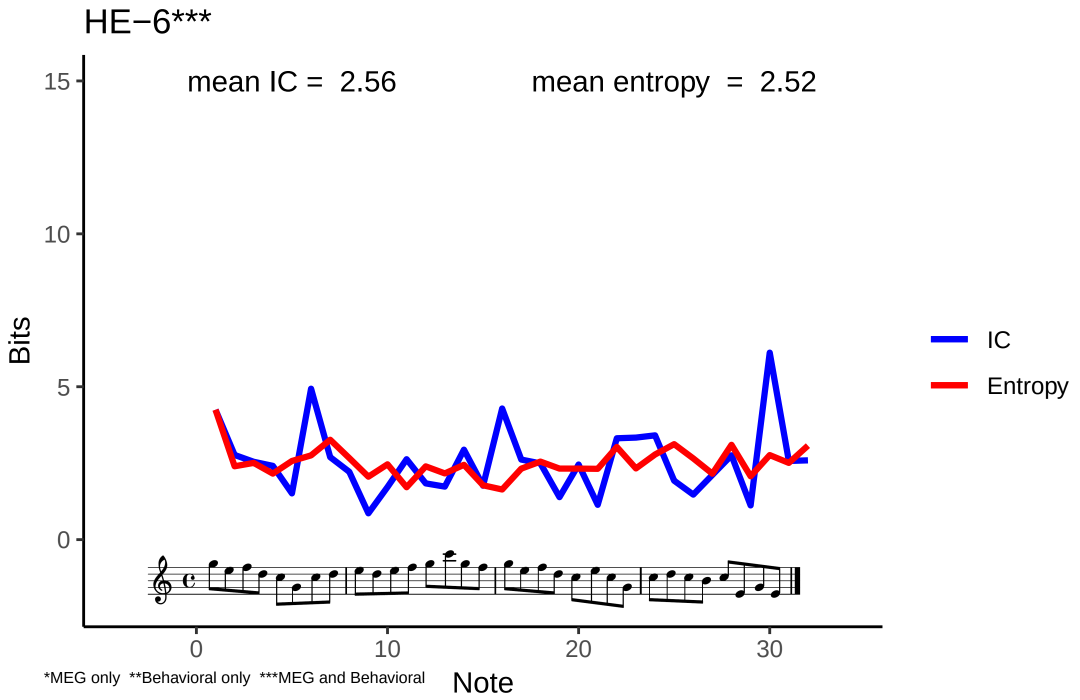
HE−6***
mean IC = 2.56
mean entropy = 2.52
Bits
Note
IC
Entropy
*MEG only **Behavioral only ***MEG and Behavioral

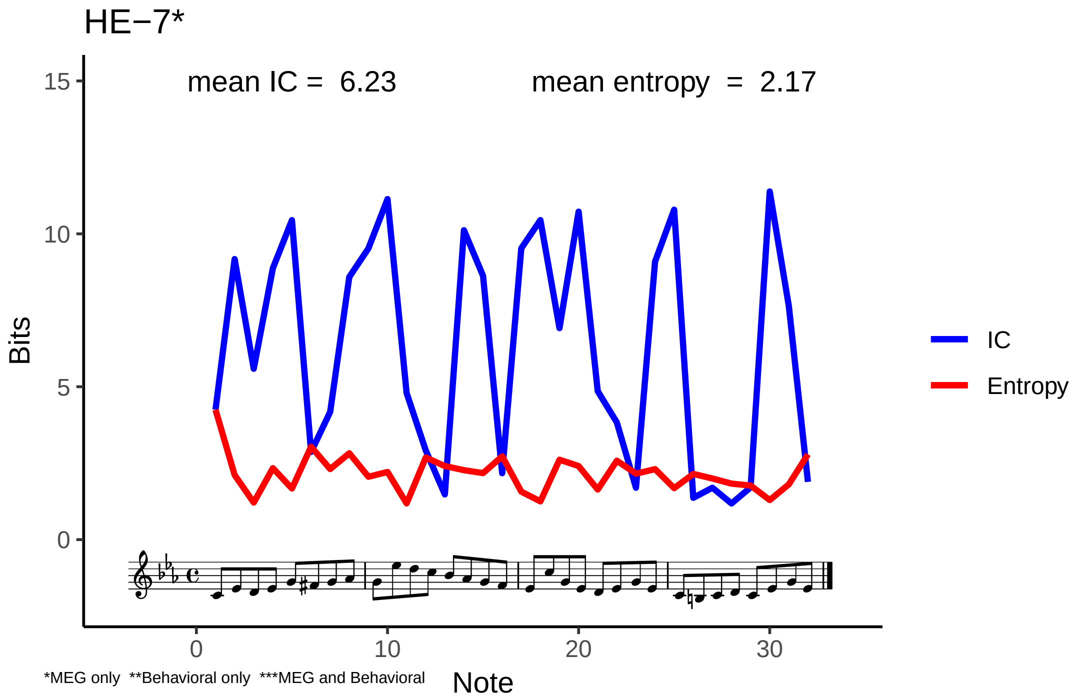

HE−7*
mean IC = 6.23
mean entropy = 2.17
Bits
15
10
5
0
0
10
20
30
Note
IC
Entropy
*MEG only **Behavioral only ***MEG and Behavioral

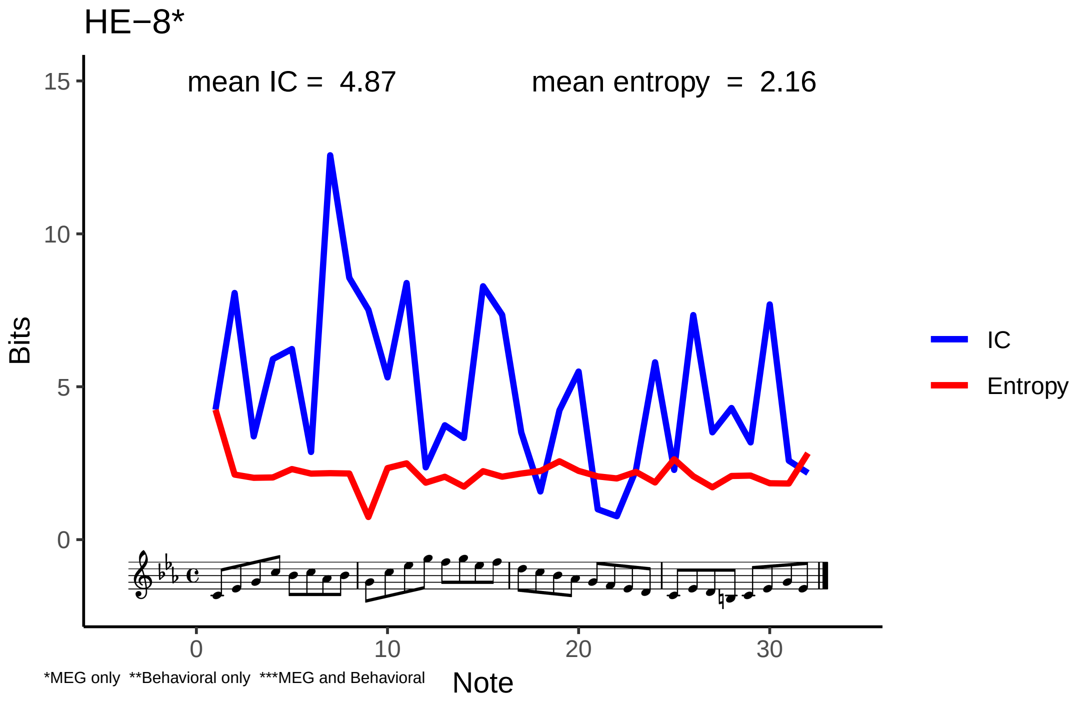

HE−8*
mean IC = 4.87
mean entropy = 2.16
Bits
Note
IC
Entropy
0
5
10
15
0
10
20
30
*MEG only **Behavioral only ***MEG and Behavioral

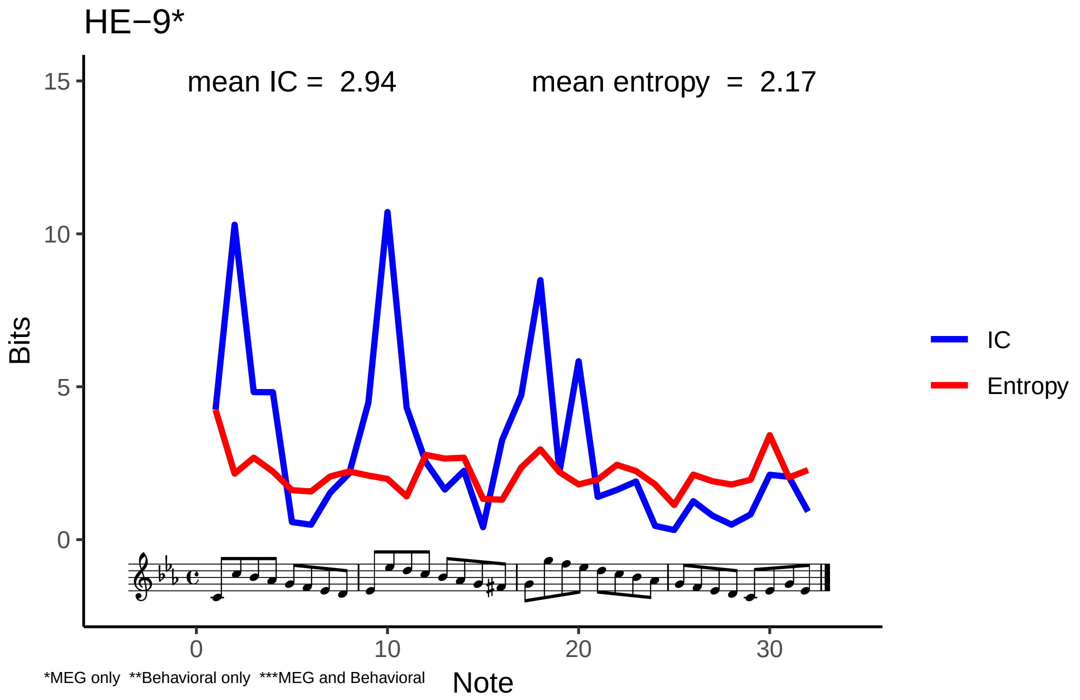

HE−9*
mean IC = 2.94
mean entropy = 2.17
Bits
15
10
5
0
0
10
20
30
Note
IC
Entropy
*MEG only **Behavioral only ***MEG and Behavioral

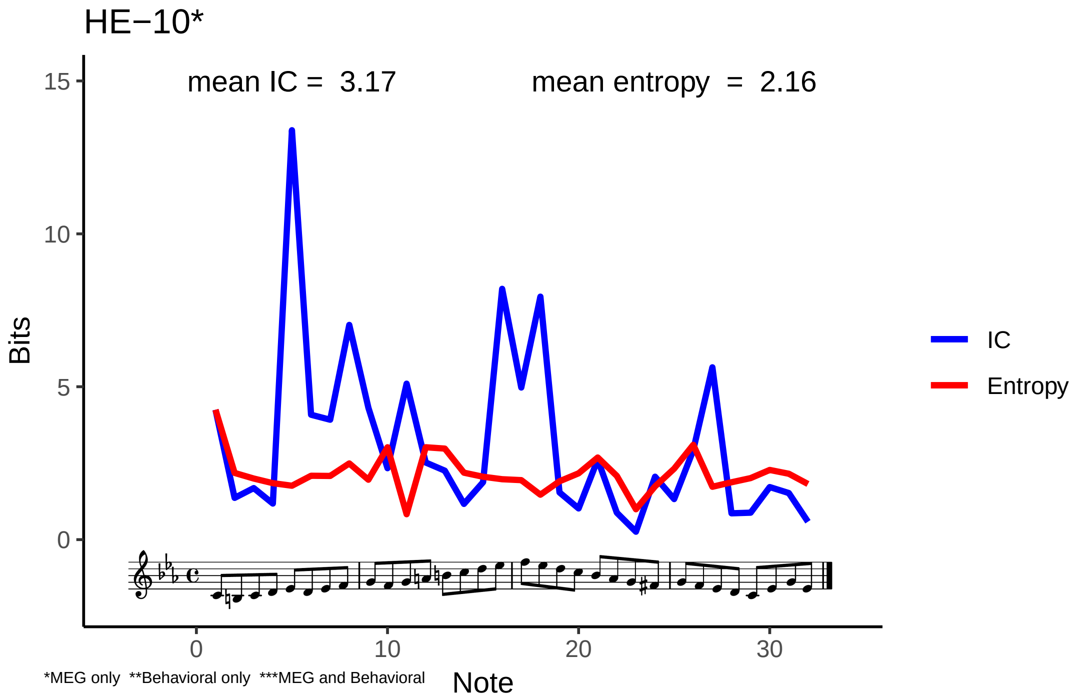
HE−10*
mean IC = 3.17
mean entropy = 2.16
Bits
15
10
5
0
0
10
20
30
Note
IC
Entropy
*MEG only **Behavioral only ***MEG and Behavioral

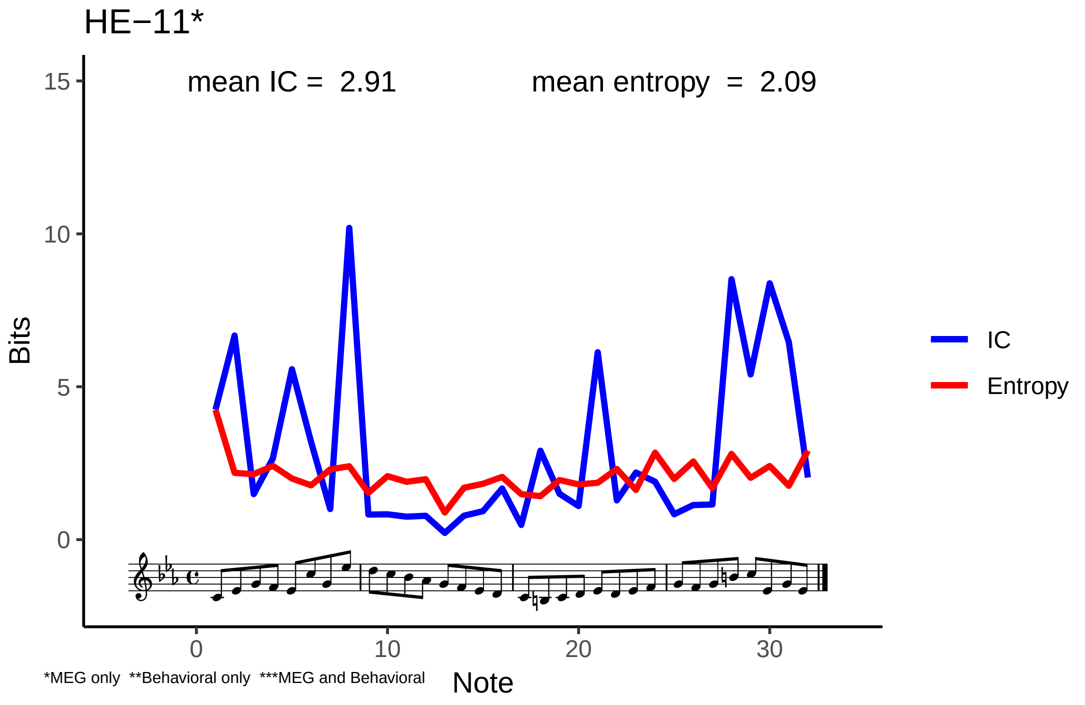
HE−11*
mean IC = 2.91
mean entropy = 2.09
Bits
15
10
5
0
0
10
20
30
Note
IC
Entropy
*MEG only **Behavioral only ***MEG and Behavioral

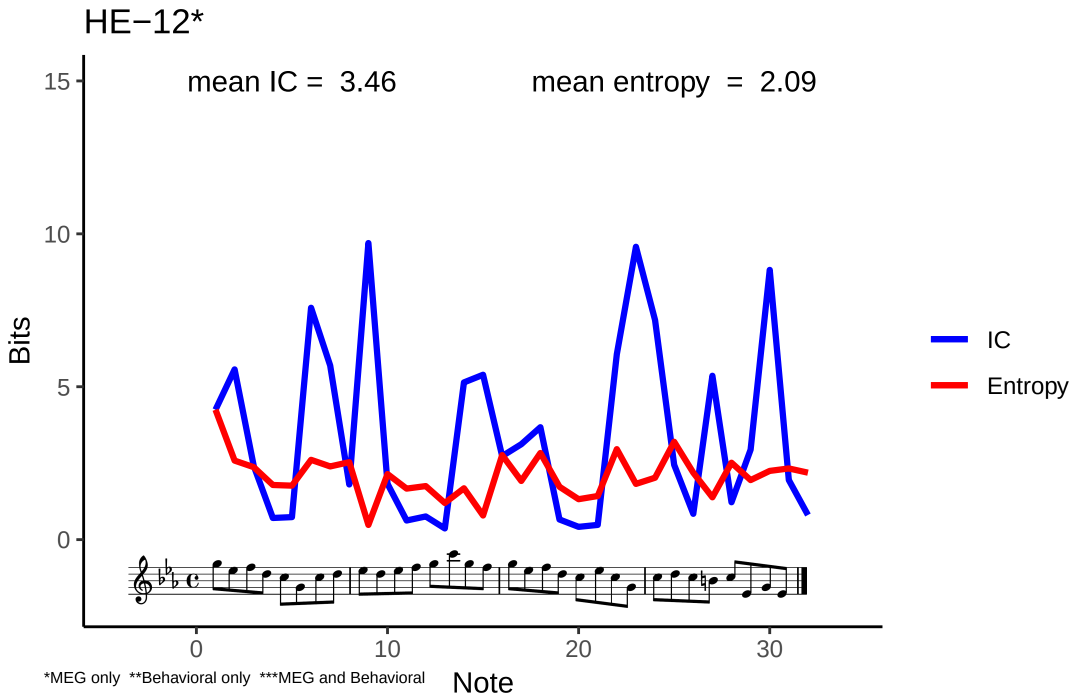
HE−12*
mean IC = 3.46
mean entropy = 2.09
Bits
15
10
5
0
0
10
20
30
Note
IC
Entropy
*MEG only **Behavioral only ***MEG and Behavioral
