## Supplementary figures and images for "Reduced prediction error responses in high- as compared to low-uncertainty musical contexts"

### Supplementary Figure 1

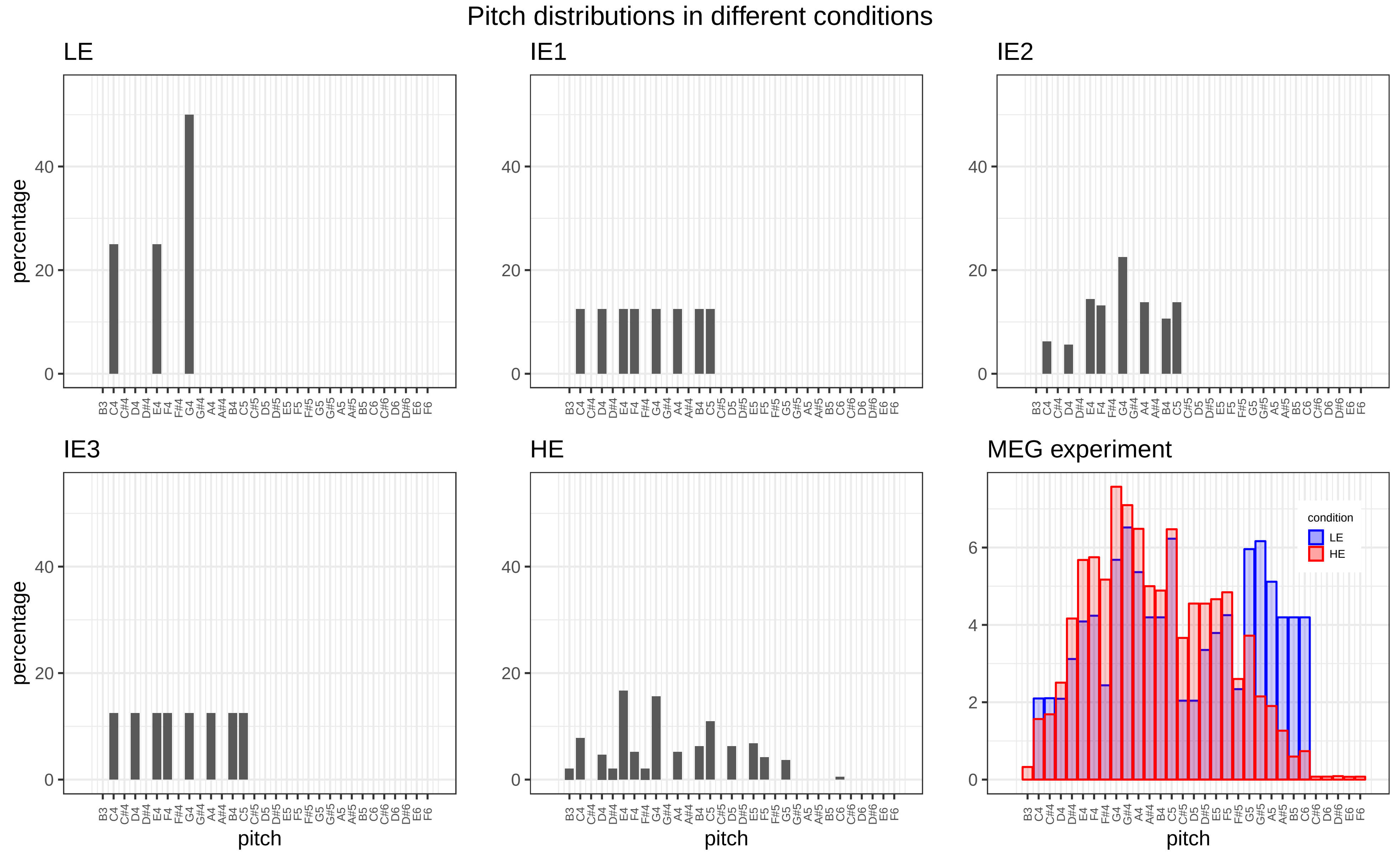

### Supplementary Figure 2

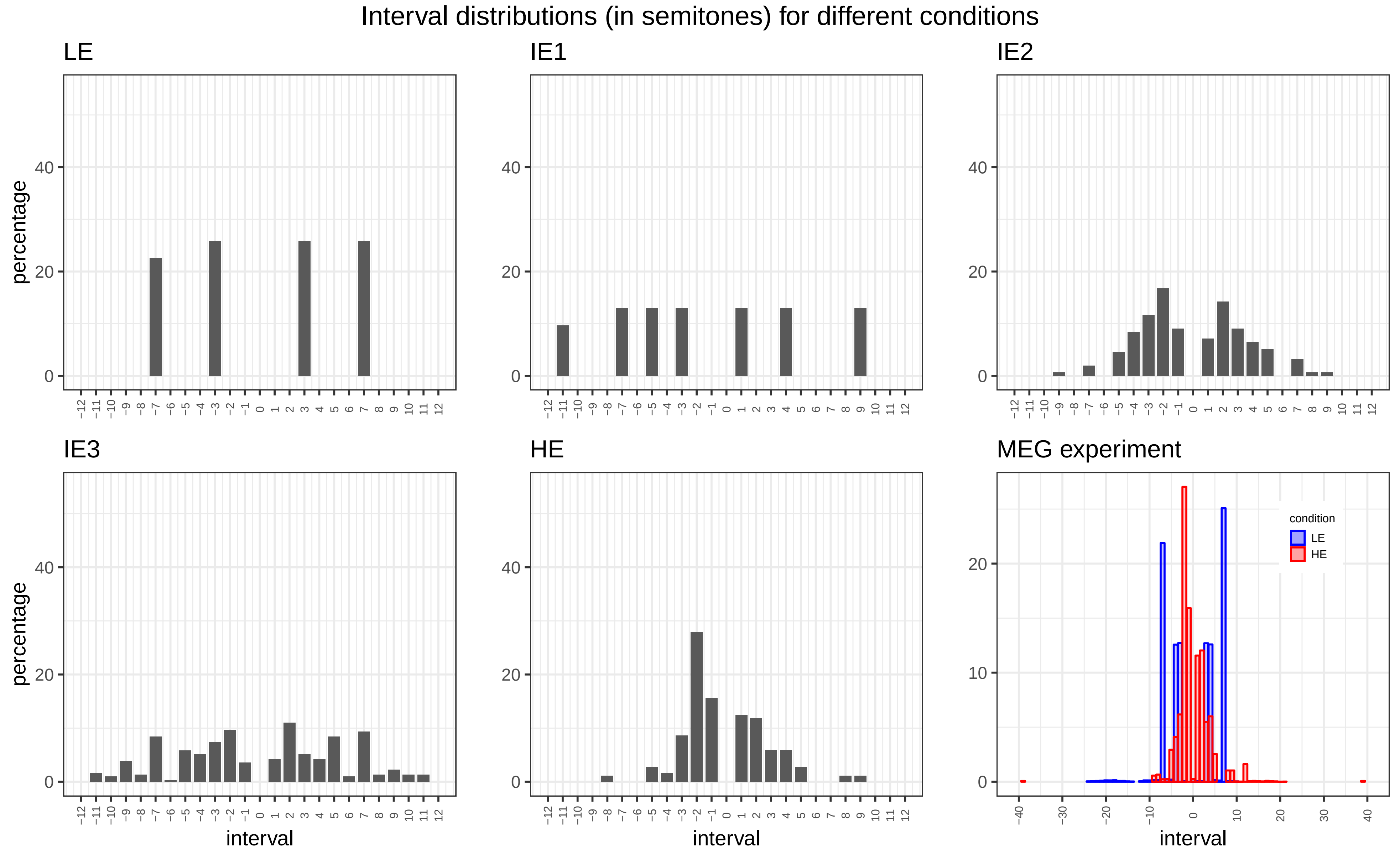

### Supplementary Figure 3

Adjusted and unadjusted IDyOM estimates for MEG (A,C) and behavioral (B,D) experiments

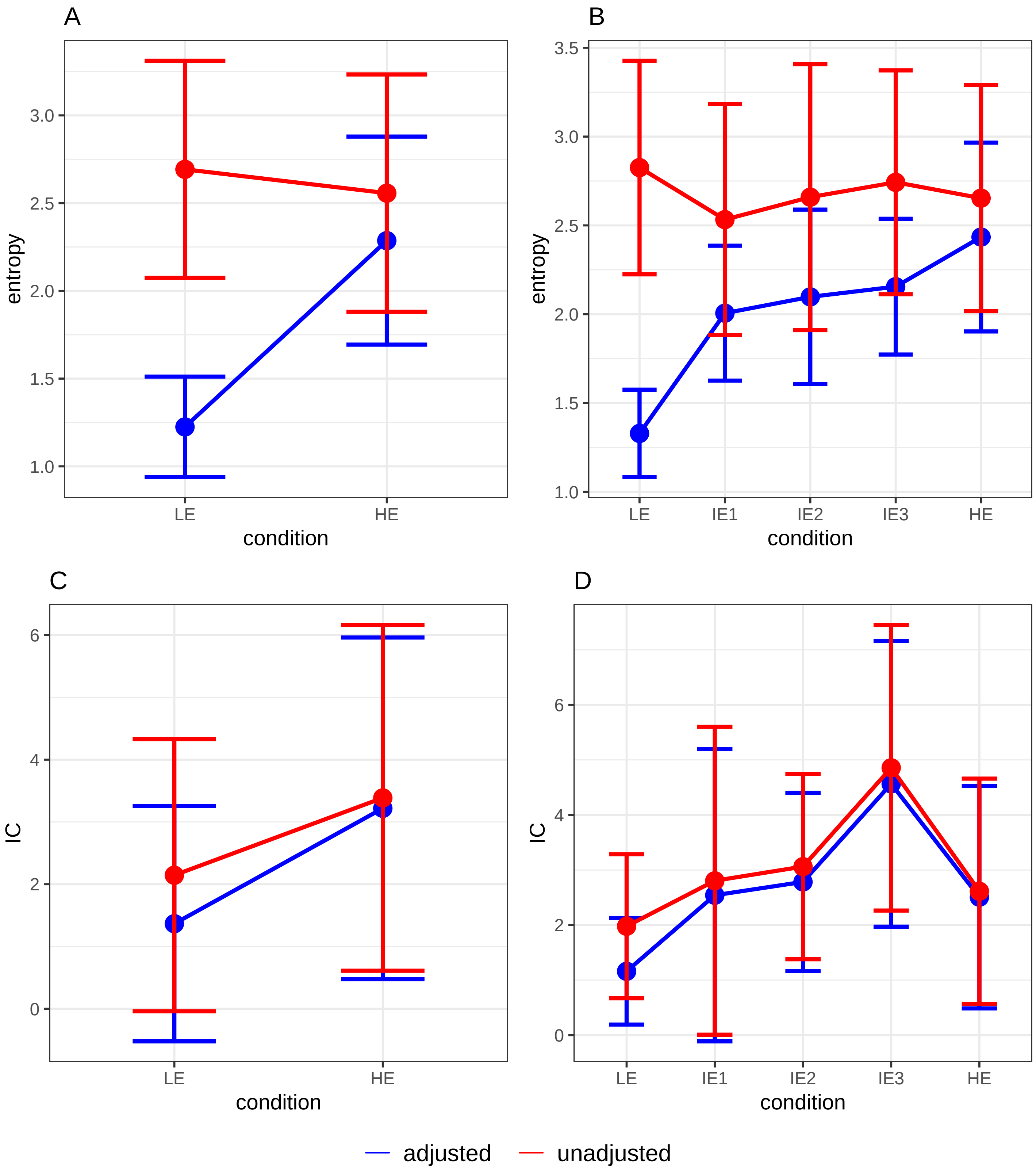

### Supplementary Figure 4

Adjusted mean IDyOM estimates for each melody in MEG (A,C) and behavioral (B,D) experiments

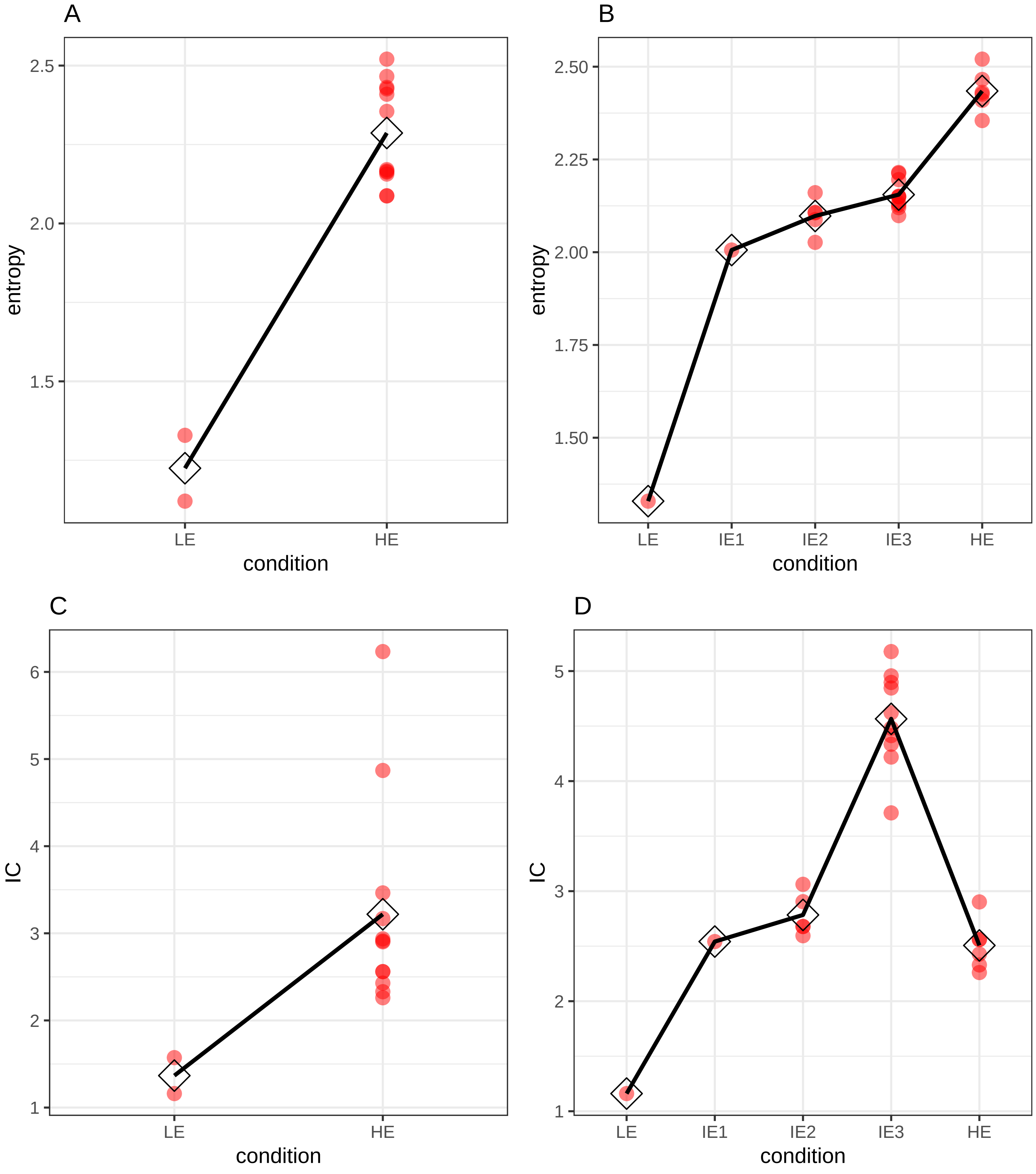
